# Surprising regulatory plasticity for the conserved HOG pathway in diverse *Saccharomyces cerevisiae* strains

**DOI:** 10.64898/2025.12.28.696518

**Authors:** Carson L. Stacy, Amanda N. Scholes, Tara N. Stuecker, Stephanie E. Hood, Crystal C. Crook, Maria Espana-Pena, Adam C. Paré, Jeffrey A. Lewis

**Affiliations:** Interdisciplinary Graduate Program in Cell and Molecular Biology, University of Arkansas, Fayetteville, Arkansas 72701, United States of America; Department of Biological Sciences, University of Arkansas, Fayetteville, Arkansas 72701, United States of America

## Abstract

Mitogen-activated protein kinases (MAPKs) display remarkable regulatory plasticity across evolution, ranging from highly specialized pathways to broadly responsive global signaling hubs. In the budding yeast *Saccharomyces cerevisiae*, the high-osmolarity glycerol (HOG) network has served as a paradigm for largely stress-specific MAPK signaling, where the Hog1 MAPK coordinates osmoadaptation. This stands in sharp contrast to Hog1 orthologs in other fungi and humans, which respond not only to osmotic stress, but also diverse stresses including UV, heat shock, oxidative stress, and pathogen signals. Whether the relative osmospecificity of *S. cerevisiae* Hog1 represents an ancestral feature or lineage-specific evolution remains unclear. The majority of foundational work on *S. cerevisiae* HOG signaling has been performed in laboratory strains that are known to be genetic and phenotypic outliers, and we have been leveraging wild *S. cerevisiae* strains to understand aspects of stress signaling that may have been lost in laboratory strains. Here, we examined the phenotypic effects of *hog1Δ* mutations in a commonly-used laboratory *S. cerevisiae* strain and diverse wild strains on stress cross protection and gene expression. Our findings demonstrate an expanded role in cross-stress protection for Hog1 in wild yeast strains compared to the laboratory strain. More strikingly, we identified a large number of Hog1-dependent genes for non-osmotic stresses in the wild strains that were completely absent in the lab strain. Notably, the Hog1 regulon in wild strains responding to non-osmotic stresses is largely distinct from the canonical osmotic stress response, which we show likely occurs through non-canonical cytoplasmic functions. These findings reveal surprising within-species plasticity for the highly conserved HOG network, suggesting that evolutionary transitions between specialist to generalist stress signaling may occur with relative ease.

## Introduction

A key feature of eukaryotic stress defense is gene expression remodeling orchestrated by diverse mitogen-activated protein kinases (MAPKs) [1]. One of the best studied MAPK pathways is the High Osmolarity Glycerol (HOG) pathway of the budding yeast *Saccharomyces cerevisiae*, which plays a key role regulating osmotic stress responses [2]. Remarkably, HOG signaling is conserved across hundreds of millions of years of evolution, with the human homologs of yeast Hog1 (the eponymous MAPK of the HOG pathway), p38 and JNK, also playing a key role in osmotic stress signaling [3–5].

However, despite this deep conservation, these kinases differ dramatically in their specificity. While *S. cerevisiae* Hog1 predominantly responds to osmotic and related stresses [2,6], human p38 and JNK respond broadly to not only osmotic stress, but also oxidative stress, UV, heat shock, and inflammatory signals [3,7–12]. These contrasting roles raise fundamental questions about how pathway specificity evolves within conserved MAPK networks.

Canonically, during osmotic stress *S. cerevisiae* Hog1 is activated via two complementary upstream sensory branches, Sln1 and Sho1, which converge on the MAPK kinase Pbs2 to ultimately phosphorylate Hog1 [6,13]. Once phosphorylated, Hog1 translocates to the nucleus within minutes, leading to dramatic transcriptional remodeling [13,14] including induction of key stress defense genes (e.g., those such as *GPD1* that increase the concentration of osmolytes such as glycerol) and repression of ribosomal protein genes thought to redirect translational capacity to induced transcripts [15–17]. In the nucleus, Hog1 directly phosphorylates osmo-specific transcription factors including Hot1, Sko1, and Smp1 [18–21], while also regulating the general stress transcription factors Msn2/4 [19], thereby upregulating hundreds of genes. Hog1 itself also directly recruits RNA Pol II to osmostress-regulated genes [22]. Beyond transcription, Hog1 has also been implicated in global regulation of translation via phosphorylation of another MAPK, Rck2 [16,23–25]. Hog1 also has cytoplasmic phosphorylation targets during osmotic stress [26,27]. For example, it directly phosphorylates and inactivates the Fps1 aquaglyceroporin to ensure that produced glycerol does not exit the cell [28] and has been shown to phosphorylate heat shock protein Hsp26 [29], leading to activation [30]. The diversity of Hog1’s regulatory targets, multiple modes of activation, and cross-talk with other stress pathways such as the cell wall integrity (CWI) pathway [31] all highlight the potential for plasticity in this highly conserved signaling network.

A major question in the evolution of the HOG network is why it is largely osmospecific in *S. cerevisiae*, but is generally stress responsive in not only multicellular organisms such as humans, but also in some pathogenic yeast species such as *Candida albicans* [32–34]. One possibility is that Hog1 is less osmospecific in *S. cerevisiae* than previously appreciated, and that the perceived osmospecificity may also be dependent on genetic background. Indeed, there has been an accumulating body of evidence suggesting that Hog1 does play some role in responding to other stresses beyond osmotic, including oxidative heat, cold, and small molecule stress responses [25,28,35–44] . However, evidence for a non-osmotic role for Hog1 has been predominantly phenotypic (fitness defects in *hog1Δ* mutants) or based on detecting phospho-Hog1 [25,39,40,45,46], with essentially no systematic interrogation of the transcriptional response to non-osmotic stress in *hog1Δ* mutants. Transcriptional effects ascribed to Hog1 for ‘non-osmotic’ stresses examined to date including extremely low pH and hypoxia [47–50] are thought to mimic hyperosmotic stress and thus activate HOG signaling via membrane alterations. Overall, the lack of systematic study of the transcriptional response of *hog1Δ* mutants to diverse stresses remains a major gap in our understanding of the presumed osmo-specificity of *S. cerevisiae* Hog1.

Additionally, the majority of studies of Hog1 function have been in domesticated laboratory strains that we now know are extreme genetic and physiological outliers [51–55]. We have been taking advantage of natural variation in wild yeast strains to identify novel features of yeast stress defense that may have been lost in these laboratory strains. In particular, we have been leveraging phenotypic variation in a phenomenon called acquired stress resistance, where cells exposed to a mild “primary” dose of stress can gain resistance to an otherwise lethal “secondary” dose of stress. Notably, acquired stress resistance can manifest as same stress protection or cross protection. We and others have found that stress-activated gene expression changes correlate better with acquired stress resistance than the basal resistance of unstressed cells [53,56]. Moreover, because naturally occurring stresses often occur in gradients or in predictable patterns of succession, acquired stress resistance is likely relevant for fitness in natural environments.

We previously found that our commonly-used laboratory yeast strain (S288C) fails to acquire further H_2_O_2_ resistance following mild ethanol treatment, while the vast majority of wild yeast strains can [53]. We subsequently found that the defective cross protection in the lab strain was due to a mutation in *HAP1* encoding a transcription factor, which led to decreased expression of a key H_2_O_2_ scavenging enzyme (Ctt1) [57]. We subsequently sought to identify other key regulators of cross protection in wild yeast strains, which led us to examine the role of Hog1 in both S288C and diverse wild yeast strains. As expected, Hog1 played a key role in cross protection for all strains when the primary stress was osmotic (NaCl). Surprisingly however, we found that Hog1 played a large role in cross protection in diverse wild strains for non-osmotic stresses as well, which was not observed in the S288C-derived lab strain. To understand whether gene expression variation was responsible, we compared the transcription responses of wild-type and *hog1Δ* mutants across strains and stresses.

While Hog1 was largely osmospecific in S288C, Hog1 played a much larger role in the response to H_2_O_2_ and ethanol in wild strains. Mechanistically, we found that HOG signaling for these non-osmotic stresses likely occurs non-canically via cytoplasmic functions rather than through canonical nuclear localization. Altogether, this study reveals surprising plasticity in HOG signaling in wild yeast strains, and has important implications for the supposed osmospecificity of Hog1 in *S. cerevisiae*.

## Materials and Methods

### Yeast strains and growth conditions

Strains and primers used in this study are listed in Supplementary Tables S1 and S2, respectively. Parental strain backgrounds include a S288C-derived laboratory strain (DBY8268), and three wild isolates: YPS606 (oak-soil), M22 (vineyard), and YJM1129 (distillery). All strains are prototrophic homozygous diploids (construction described in [58]). Deletions of *HOG1* in all strain backgrounds were constructed by PCR amplifying and transforming the *hog1Δ*::KanMX cassette from the Yeast Knockout Collection [59] into each parental strain, followed by sporulation and tetrad dissection as described to obtain homozygous deletions (all strains are homothallic and thus capable of mating-type switching and selfing) [60]. All deletions were verified via diagnostic PCR with primer pairs that anneal to the MX cassette and a sequence upstream of the deletion (verifying the ‘scar’), and primer pairs that anneal within the deleted region (verifying gene absence). To generate heterozygous C-terminal Hog1-GFP fusions, GFP was amplified from pFA6a-GFP(S65T)-NatMX6 (Euroscarf P30416) [61] and transformed into the appropriate wild-type strains. Diagnostic PCR using primers that amplify the *HOG1-GFP* junction were used to confirm construction of heterozygous *HOG1*-GFP fusion strains [62].

All yeast strains were grown at 30°C with orbital shaking (270 rpm) in YPD (1% yeast extract, 2% peptone, 2% dextrose) or synthetic complete (SC) [63]. Sporulation was induced by growing cells for 48 h in YPD, collecting cells by gentle centrifugation (1,500 x *g* for 3 min), resuspension in 1% potassium acetate, and incubating for 2-10 days at 25°C with 270 rpm orbital shaking. Yeast transformations were performed via the high-efficiency method of [64]. When used for selection, antibiotic concentrations were 100 µg/ml for nourseothricin (Nat) and 200 µg/ml for G418 (Kan).

For growth curve analysis of *HOG1* mutants in mild stress conditions, cells were grown overnight in YPD with orbital shaking (270 rpm) at 30°C to saturation on YPD. Cells were then diluted into 1 mL of YDP media in 24-well plates at an OD_600_ of 0.1, with the following stress concentrations: 0.4 mM H_2_O_2_, 5% ethanol, or 0.4M NaCl. Plates were sealed with a Breathe Easy sealing membrane (Electron Microscopy Supply), and growth was monitored every 15 min over 24 h in an Eon microplate spectrophotometer (BioTek) with continuous orbital shaking at 30°C. Growth experiments were performed in biological triplicate.

### Cross protection assays

Cross protection assays were performed as described in [57], in biological triplicate unless otherwise noted, and a detailed protocol is available on <u>protocols.io</u> under doi doi.org/10.17504/protocols.io.g7sbzne. Briefly, ∼5 unique colonies per replicate experiment were grown overnight to saturation in YPD. Cells were then sub-cultured for at least 8 generations (∼16 hours) to mid-log phase (OD_600_ 0.3 - 0.6 on a Unico spectrophotometer). The cultures were then split, with one sample receiving a mock treatment (fresh media) and the other sample receiving a mild ‘primary’ stress treatment, with mild doses being empirically defined previously as increasing severe stress survival without affecting viability (>95% survival) [58]. Both mock and primary stress pretreatments were diluted 1:1 in fresh media, with the final concentrations of the primary stresses being 0.4 M NaCl, 0.4 mM H_2_O_2_, or 5% (v/v) ethanol. For mild heat pretreatment, cells were resuspended 1:1 in YPD prewarmed to 55°C, which immediately brought the temperature to 37°C. Mock or primary-stress pretreated cells were incubated for 1 h at 30°C (with the exception of the heat shock pretreatment which was 37°C) with 270 rpm orbital shaking, and then collected by mild centrifugation at 1,500 x *g* for 3 min. Cells were then resuspended in fresh media to an OD_600_ 0.6, and then diluted threefold into a 96-well microtiter plate containing a no-stress control and 11 severe ‘secondary’ H_2_O_2_ doses (0.75 mM, 1 mM, 1.25 mM, 1.5 mM, 1.75 mM, 2 mM, 2.5 mM, 3 mM, 3.5 mM, 4 mM, 5mM). Plates were sealed with breathable Rayon films (VWR) and incubated for 2 h at 30°C with 800 rpm shaking in a VWR Symphony shaking incubator. Cells were then diluted 50-fold in YPD in a 96-well plate, and 4 µl of each well was spotted onto YPD agar plates and grown for 48 h at 30°C before imaging.

Viability was scored semi-quantitatively using an ordinal scale of 0-4, where 0 indicated no visible colonies (fewer than 3 colonies), 1 indicated minimal growth (∼3-10 colonies), 2 indicated moderate growth (∼11-25 colonies), 3 indicated non-confluent growth, but too numerous to count (>25 colonies), and 4 indicated confluent growth (i.e., a lawn). For some experiments, viability continued past the highest secondary H_2_O_2_ concentration used initially. In those cases, an additional set of biological triplicates was performed at higher secondary concentrations to better estimate the maximum dose survived. Blinded scoring was performed using BLISS [65], with each spot being scored twice to assess scoring reliability. Scoring reliability was assessed using Cohen’s Weighted Kappa (κ) statistic with Fleiss-Cohen weighting (Cohen, J. (1960)) using the vcd package (version 1.4-13) in R version 4.4.1. The scoring reliability in this assay was “almost perfect” [66] with a Cohen’s Weighted Kappa κ=0.951 (95% CI: [0.940, 0.962]), indicating a high degree of agreement in score assignments.

Proportional-odds ordinal regression was used to estimate minimum inhibitory concentrations (MIC*), defined as H_2_O_2_ concentration where the probability of observing no viability (i.e., a score = 0) equals 0.5. MIC* analysis was performed using the ordinalMIC R package [65], with uncertainty reported as 95% confidence intervals.

### RNA extraction, library preparation, and sequencing

#### Sample Collection

RNA-seq was performed on isogenic diploid wild-type and *hog1Δ* strains in the YPS606, M22, and S288C (DBY8268) genetic backgrounds. Experiments in the YPS606 and M22 were performed in biological quadruplicate. Exploratory analysis revealed that the original S288C *hog1Δ* strain was haploid; therefore, a second set of biological triplicates was collected for S288C wild-type vs *hog1Δ* comparison using a verified diploid strain to avoid confounding ploidy effects.

Cells were grown for at least 8 generations in YPD to mid-exponential phase (OD_600_ 0.3 - 0.6).

For each biological replicate, an unstressed control sample was collected, and then the culture was split and exposed to stress media with final concentrations: 0.4 M NaCl, 5% (v/v) ethanol, or 0.4 mM H_2_O_2_. Cells were incubated in the stress for either 30 min (ethanol and H_2_O_2_) or 45 min (NaCl), timepoints which encompass the peak response for each stressor [67] and [53]. Unstressed and stressed samples were collected by centrifugation at 1,500 x *g* for 1 min, flash frozen in liquid nitrogen, and stored at -80 °C until RNA extraction.

#### RNA Extraction and Library Preparation

RNA was extracted using the ‘hot phenol’ method as described [68], with a detailed protocol available on protocols.io DOI <u>10.17504/protocols.io.inwcdfe</u>. Phenol-extracted RNA was treated with DNaseI (Ambion), and then purified with a Quick-RNA MiniPrep Plus Kit (Zymo Research), including the optional on-column DNase step. RNA quantity was measured using a Qubit fluorometer (Fisher), and all samples were confirmed to have a RIN > 9 as assessed by an Agilent 2200 Tapestation.

#### Sequencing

RNA-seq libraries were prepared using the KAPA mRNA HyperPrep Kit (Roche) and KAPA Single-Indexed Adapter Set A + B, according to manufacturer specifications with minor modifications. Starting samples consisted of 500 ng, which were then polyA-enriched, fragmented to an average size of 200-300 nucleotides, and then cDNA libraries were amplified for 9 PCR cycles. A detailed protocol is available on <u>protocols.io</u> DOI <u>10.17504/protocols.io.uueewte</u>. Libraries were pooled (24-plex), and sequenced on an Illumina HiSeq 4000 with 50 bp single-end reads at the University of Chicago Genomics Facility. To minimize potential batch effects, a random block design was used for library construction, and each biological replicate was sequenced together on a single lane. When we discovered post-analysis that the S288C *hog1Δ* strain was haploid instead of diploid, we repeated the experiment with the correct diploid strain in biological triplicate and sequenced the samples on a single HiSeq 4000 lane at Northwestern University (the University of Chicago had at that point phased out its HiSeq 4000). All raw RNA-seq data are available through the National Institutes of Health Gene Expression Omnibus (GEO) database under accession number GSE312455.

### RNA-seq analysis

#### Read Processing and Quantification

Raw reads were trimmed and filtered using TrimGalore (version 0.6.7), a wrapper for cutadapt (version 3.4), with default settings to remove Illumina adapters. Reads with phred quality scores <20 or a post-trimming length <20 bp were discarded. Trimmed reads were mapped to the S288C genome (SacCer3 R64-1-1, Ensembl accession GCA_000146045.2) using STAR (version 2.7.10a) [69] implemented through the *nf-core*/rnaseq (v 3.10.1) pipeline [70]. Alignment used a GTF containing only chromosomal protein-coding genes. Gene counts were generated using RSEM (version 1.3.1, [71]). A median of approximately 17 million reads (range: 12.4-69.5 million) were successfully mapped per sample, representing over 95% of trimmed reads successfully mapped.

#### Differential expression analysis

Differential expression analysis was performed in edgeR (v. 4.4.2) with a negative binomial quasi-likelihood (QL) generalized linear model [72,73], following TMM normalization. Genes with fewer than 10 counts across all samples were excluded from subsequent analysis. The QL model included sample type (e.g., M22 *hog1Δ* unstressed, M22 *hog1Δ* NaCl stressed, M22 *hog1Δ* ethanol stressed…) and biological replicate as factors. Contrasts of interest included comparisons of wild-type vs *hog1*Δ mutants and the stress response (stress vs. unstressed) for each strain individually (e.g., the S288C wild-type NaCl response, the S288c *hog1Δ* NaCl response), as well as comparisons of strain stress responses (e.g., S288C wild-type NaCl response vs the S288c *hog1Δ* NaCl response). Genes were considered differentially expressed if they met a Benjamini-Hochberg false discovery rate (FDR) cutoff of < 0.01 and at least a ± 1.5-fold change in expression. To identify Hog1-affected genes across all strains, all replicates were used. To normalize statistical power for between-strain comparisons, counts for each sample were subsampled down to the lowest read depth of any sample, and samples were then subset to biological triplicate.

#### Enrichment Analyses

Functional enrichment of gene ontology (GO) and Kyoto encyclopedia of genes and genomes (KEGG) categories was performed using gene set enrichment analysis (GSEA) using the compareCluster function in clusterProfiler (version 4.14.4) [74], with significance assessed by Fisher’s exact test (FDR < 0.01). GSEA analysis utilizes estimated log fold change estimates for all transcripts in the transcriptome, regardless of significance cutoffs or number of genes identified as differentially expressed. Enrichment for transcription factor (TF) targets was identified by GSEA against small and large-scale linkage evidence from the TFLink database [75].

#### Phenotype-Expression Correlation Analysis

Partial least squares regression (PLS1) was used to identify correlations between gene expression changes and cross-protection phenotypes (H_2_O_2_ MIC*). Log-normalized counts for genes were used as predictors, and MIC* served as the response variable. To retain the importance of large expression changes, log-normalized counts were not further rescaled. Because larger changes in log count per million values are likely more important than smaller changes, predictors were not rescaled to a variance of unity for each gene. PLS1 was chosen for its ability to identify associations in high-dimensional data [76–78]. Alternative approaches, including elastic net, ridge and lasso regression, principal component regression, extreme gradient boosting, and random forest machine learning algorithms were also tested for phenotype correlation; however, PLS was chosen because it predicted the known true positive *CTT1* as among the top targets in preliminary analyses. The model was fitted using the plsr function from the pls R package (v. 2.8-5) [76]. Because the phenotyping and RNA-seq experiments were not paired (i.e., replicate 1 of a cross protection experiment did not correspond to the same cells as replicate 1 of an RNA-seq experiment), we used a bootstrapping approach (N=1000 resamples) to randomly pair phenotype and expression replicates within each strain-genotype-pretreatment condition. This approach avoids potential spurious correlations that could arise from artificial pairings, while preserving the biological structure of the data. The number of PLS components was optimized in each bootstrap via 10-fold cross-validation. Variance Inflation Factor (VIF) estimates were calculated for each gene, with a standard VIF cutoff of 1.

### Hog1 phospho-westerns

Hog1 phosphorylation was assessed in biological duplicates using heterozygous *HOG1*-GFP::NatMX diploids in the S288C, M22, and YPS606 strain backgrounds. Cells were grown to mid-exponential phase and treated with mock (media alone), 0.4 M NaCl, 0.4 mM H_2_O_2_, or 5% (v/v) ethanol. Samples were incubated at 30°C with 270 rpm orbital shaking, and then collected at 5, 30, and 60 min post-stress. Cell pellets were flash frozen in liquid nitrogen and stored at -80°C until processing.

Samples were processed as described [79,80], with minor modifications. Briefly, samples were resuspended in different volumes of lysis buffer (0.1M NaOH, 50m EDTA, 2% SDS, 2% β-mercaptoethanol and protease inhibitor tablets [VWR A32963]) to achieve identical cell concentrations across samples (∼1 x 10^7^). Two-hundred µl of each sample was then lysed by incubating at 95°C for 10 min, followed by the addition of 5 µl of 4 M acetic acid and vortexing for 30 sec, and then followed by a second incubation at 95°C for 10 min. Fifty µl of Laemmli buffer was added to each sample, and cellular debris was pelleted via microcentrifugation at 21,130 x *g* for 5 min. Following SDS-PAGE, proteins were transferred at 4°C for 1 h onto PVDF membranes (VWR catalog number 10120-024). The membrane was blocked for 1 h with 5% milk in Tris Buffered Saline (TBS) with gentle rocking, and primary antibody incubation was performed at 4°C overnight in 5% milk in TBST (0.1% Tween 20).

Phospho-Hog1 was detected using mouse anti-phospho-p38 MAPK (Thr180/Tyr182) (28B10) (Cell Signaling catalog number 9216) at a 1:2000 dilution, and actin (loading control) was detected via rabbit anti-actin (Invitrogen catalog number PIPA585271) at a 1:4000 dilution. The secondary antibody incubation was done at room temperature using a dilution of 1:10,000 for 1 h with gentle agitation in 5% milk in TBST-SDS (0.1% Tween 20 and 0.01% SDS), using IRDye 800CW donkey anti-mouse (Li-Cor Biosciences catalog number 926-32212) and IRDye 880RD donkey anti-rabbit (Li-Cor Biosciences catalog number 926-68073). Blots were imaged using a Li-Cor Odyssey scanner using Image Studio v2.0. Brightness and contrast were adjusted globally, and no bands were lost or obscured with adjustments. Raw western blot images can be found on Zenodo doi: 10.5281/zenodo.18050221.

### Hog1-GFP microscopy

Hog1-GFP localization experiments were performed in biological triplicate using heterozygous *HOG1*-GFP::NatMX diploids. Cells were grown >8 generations in SC at 30°C with 270 rpm orbital shaking to mid-exponential phase (OD_600_ of 0.3-0.6). Once cells had reached mid-exponential phase, DAPI was added to a final concentration of 2.5 µg / ml, and cells were returned to the incubator for 30 min. Cells were collected by centrifugation at 1,500 *g* x 1 min, and pellets were resuspended in 100 µl of either mock treatment (SC alone) or final concentrations of stress treatments of 0.4 M NaCl, 0.4 mM H_2_O_2_, or 5% (v/v) ethanol, with 2.5 µg / ml DAPI maintained throughout treatments. Live imagining was performed at exactly 4 min of stress exposure, as well as 5-min time-windows from 5 to 30 minutes, and 60 min for YPS606 in H_2_O_2_ stress, using an Axio Observer 7, Zeiss LSM900 confocal microscope with a Plan-Apochromat Neo20x/0.8 NA dry objective. Images were collected using the ZEN 3.0 software program (Zeiss). Uniform brightness and contrast adjustments were performed using Fiji package [81]. Raw microscopy images can be found on Zenodo doi: 10.5281/zenodo.18050221.

### Data and Code Availability

All raw RNA-seq data generated in this study are available at the NCBI Gene Expression Omnibus (GEO) under accession number GSE312455. The *S. cerevisiae* reference genome SacCer3 R64-1-1 and annotation were obtained from Ensembl Release 104. All analysis code is available on Github: https://github.com/LewisLabUARK/Stacy2026_HOG1, and raw data for figures are available on Zenodo: doi: 10.5281/zenodo.18050221

## Results

### Hog1 is required for non-osmotic cross protection in wild and lab *S. cerevisiae* strains

We previously found that an S288C-derived laboratory strain of yeast was unable to acquire further H_2_O_2_ resistance following pretreatment with mild ethanol stress [57], which was due to defective induction of a key H_2_O_2_ scavenger (*CTT1* encoding catalase). To identify possible regulators controlling stress cross protection in wild yeast strains, we performed targeted mutagenesis of candidate regulatory genes implicated by transcriptional profiling [58]. We ultimately found conditional specificity, where in a wild oak strain (YPS606) the paralogous general stress defense transcription factors Msn2/4 are necessary for ethanol-induced, but not salt-induced, cross protection against H_2_O_2_.

Because the stress-activated MAP kinase Hog1 is a known regulator of Msn2/4 during osmotic stress [82], and is required for NaCl-induced cross protection against H_2_O_2_in S288C [56], we included *hog1*Δ mutants in both the S288C and YPS606 backgrounds as ‘positive’ controls for acquired stress resistance assays. As expected for S288C, the *hog1*Δ mutant had defective acquired H_2_O_2_resistance following NaCl pretreatment, and little detectable acquisition following ethanol pretreatment (Fig. 1A). Surprisingly however, the YPS606 *hog1*Δ mutant retained substantial NaCl-induced cross protection against H_2_O_2_, but displayed a clear defect when ethanol was the primary stressor (Fig. 1A). To better quantitate strain-specific differences for the role of *HOG1* in cross protection against H_2_O_2_, we included *hog1*Δ mutants in a wild vineyard strain (M22) and a distillery strain (YJM1129). We additionally increased the maximum doses of H_2_O_2_, allowing us to better estimate the minimal inhibitory concentration (MIC*) of H_2_O_2_ for each experiment (see Methods). As expected, NaCl pretreatment robustly cross protected against severe H_2_O_2_ survival in all strain backgrounds, with about half of the level of cross-protection dependent on *HOG1* (Fig. 1B). Similar to previous findings [57,58], S288C did not acquire H_2_O_2_ resistance following ethanol pretreatment. Notably though, S288C also displayed lower levels of NaCl-induced H_2_O_2_ resistance than any of the other strains tested (MIC* = 7.53 vs a range of 12.6-33.6 for the wild strains; Table S3). Surprisingly, *HOG1* was required for cross-protection in at least one non-osmotic treatment in at least one wild strain (Fig. 1B, Fig. S1, and Table S4). For example, *HOG1* had large effects on acquired H_2_O_2_ resistance when H_2_O_2_ was the pretreatment for YPS606 (ΔMIC* = -3.04, p-value = 1.8×10^-9^), when ethanol was the pretreatment in M22 (ΔMIC* = - 1.61, p-value = 1.34×10^-4^), and when heat was the pretreatment in YPS606 (ΔMIC* = -2.35, p-value = 2.8×10^-6^). Altogether, these results suggest that Hog1 is important for acquired H_2_O_2_resistance not only following osmotic stress pretreatment, but also following mild oxidative, ethanol, and heat stresses.

**Figure 1.**
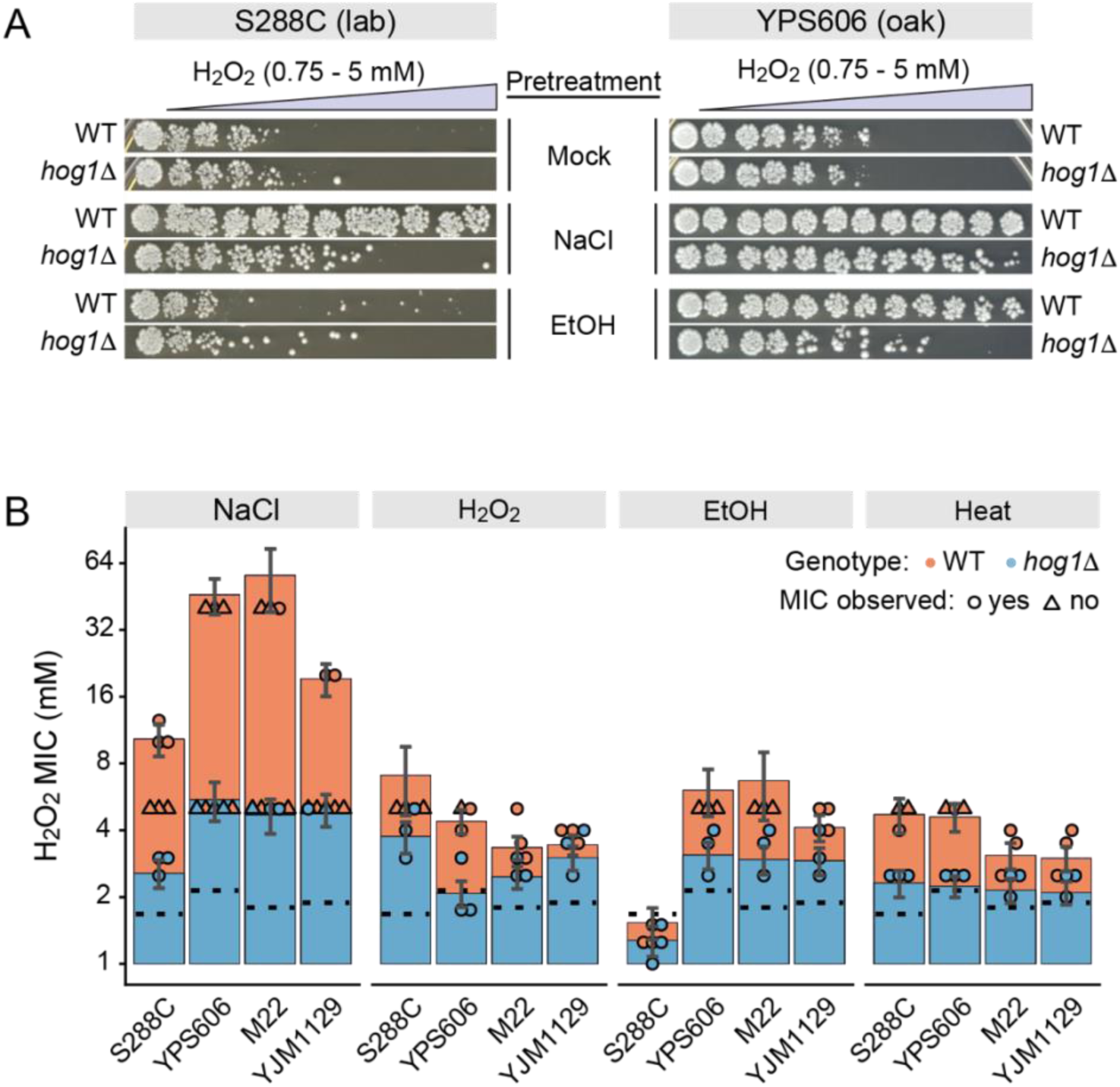
Unexpected diversity in the conditional requirement for *HOG1* in stress cross protection across yeast strains. (A) Representative H_2_O_2_ cross protection assays for S288C and YPS606. Cells were exposed to either 5% (v/v) ethanol, 0.4 M NaCl, or mock (water) pretreatments for 60 min, the mild stress was removed, and then cells were exposed to 11 doses of severe H_2_O_2_ for 2 h. Cells were then plated to score viability. (B) Blinded scoring software (BLISS) was used to generate ordinal scores, which were then used to calculate MIC*, which is the estimated concentration of H_2_O_2_ at which the probability of no viability (i.e., a score of 0), exceeds 0.5 (see Methods). Error bars denote 99% confidence intervals for MIC*. Individual data points for each replicate indicate either the observed MIC for a replicate (circle), or the maximum concentration survived when MIC was not directly observed (triangle). Black dotted lines denote the peroxide resistance of each strain without pretreatment.

### Hog1 regulates the transcriptional response to non-osmotic stresses in wild yeast strains

That Hog1 was required for cross-stress protection following diverse pretreatments in wild strains, but not S288C, raised the question of whether HOG signaling is less osmospecific in natural populations. To test this, we performed transcriptional profiling of both wild-type and *hog1*Δ mutants S288C and the two wild strain backgrounds where Hog1 had the largest effects on non-osmotic stress cross protection phenotypes (M22 and YPS606). We compared expression of unstressed cells, as well as cells exposed to 0.4M NaCl, 0.4mM H_2_O_2_, or 5% (v/v) ethanol stress.

As expected, for all strains the largest number of differentially expressed genes between the WT vs *hog1*Δ strains were observed for the NaCl response (FDR < 0.01 and fold change > 1.5; Fig. 2). Notably, the wild strains did have higher numbers of Hog1-dependent genes for the NaCl response compared to S288C (1,394 in YPS606, 915 in M22, and 251 in S288C; Table S5). Additionally, the ‘directionality’ of the Hog1 effect also differed in S288C versus the wild strains, with S288C *hog1Δ* mutant showing more cases of defective repression, and the *hog1Δ* mutants in the two wild strains showing enhanced repression. The wild strain *hog1Δ* mutants also exhibited more defective induction of salt-responsive genes. Altogether, the number of differentially expressed genes and their directionality in the wild-type vs *hog1Δ* strain comparisons suggest that HOG signaling both differs and is reduced in S288C for the osmotic stress condition tested. To directly ensure that these observed differences across strains were not due to lack of power to detect differential expression in S288C compared to the wild strains, we randomly downsampled all libraries to the same minimum depth (∼12 million reads) and replicate numbers (biological triplicates). The overall trend remained unchanged, with S288C having both fewer Hog1-regulated genes during NaCl stress and different directionality (Fig. S2). Wild-type S288C also had fewer differentially expressed genes during NaCl stress compared to the wild strains (937 in S288C vs 2404 inYPS606 and 2782 in M22; Fig. S3), which we also observed previously [58]. This observation suggests that S288C differs from the wild strains in either osmosensing or stress perception, potentially due to variation in even canonical HOG signaling (see Discussion).

**Figure 2.**
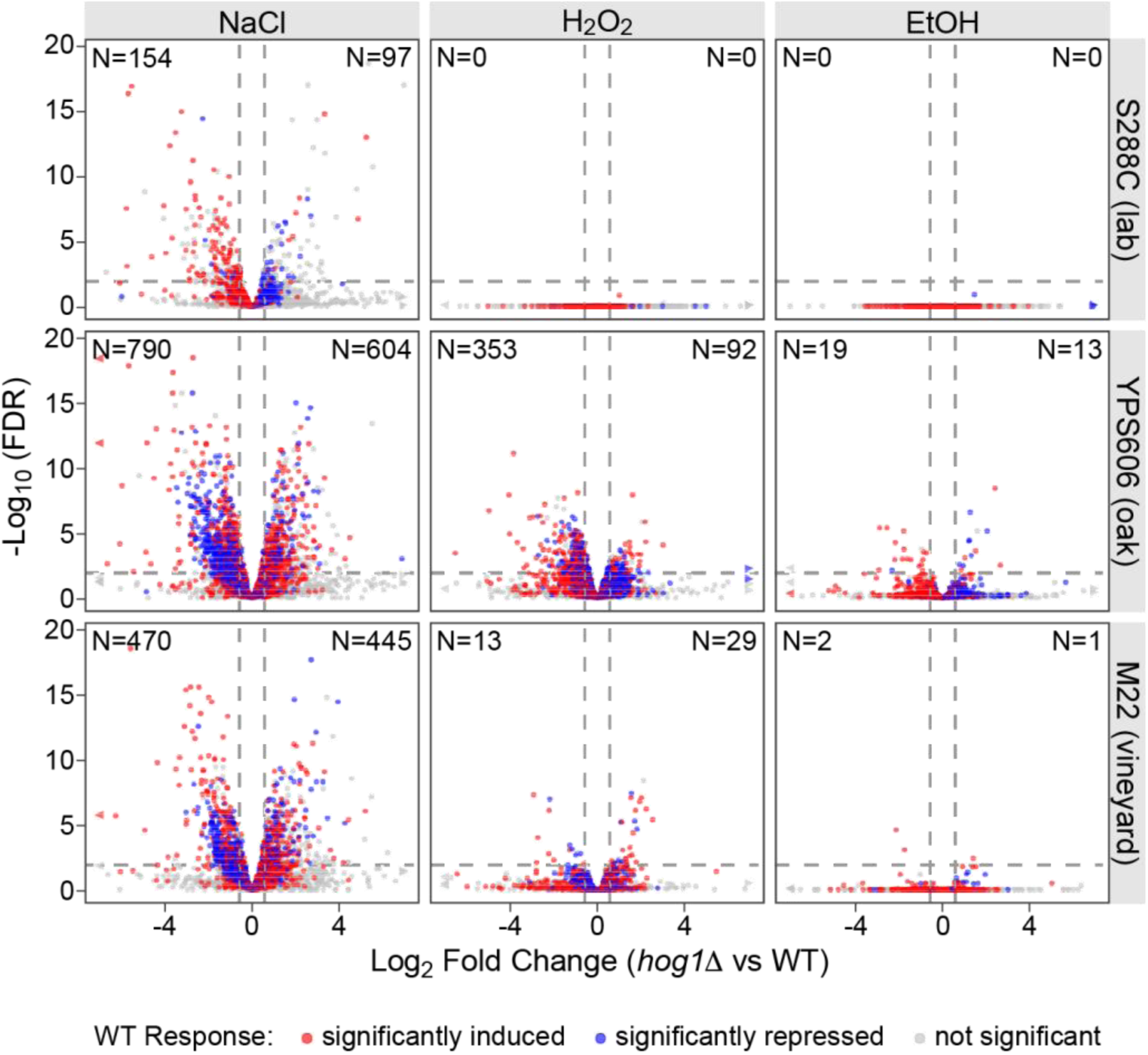
*HOG1*-dependent transcriptional changes are osmospecific in S288C but extend to non-osmotic stresses in wild strains. Volcano plots depict Hog1-dependent gene expression changes during the depicted stresses (columns) for each strain (row). Each point represents a gene, and for each sub-panel, genes with significantly reduced stress-responsive expression in the *hog1Δ* mutant vs the wild-type strain are in the upper-left quadrant (negative log_2_ fold change values), while genes with significantly higher expression in the *hog1Δ* mutant vs the wild-type strain are in the upper-right quadrant (positive log_2_ fold change values). Point colors indicate each gene’s behavior in wild-type cells responding to the depicted stress: red indicates genes significantly induced (>1.5-fold, FDR < 0.01), blue indicates a gene significantly repressed, and gray indicates genes with no significant change. (N) denotes the number of Hog1-dependent genes in each direction within each panel.

We also notably observed substantial numbers of Hog1-dependent genes during non-osmotic stresses in the two wild strains, but not in S288C. In YPS606, hundreds of genes were dependent on Hog1 for the H_2_O_2_ response (Fig. 2), and dozens were Hog1-dependent during ethanol stress. The effect of Hog1 was not as strong in M22, but there were still dozens of Hog1-regulated genes for the H_2_O_2_ response, and a small number of the ethanol response. In contrast, we observed zero Hog1-dependent genes in S288C for either the H_2_O_2_ or ethanol responses. Again, the inability to detect significant Hog1-regulated genes during the S288C H_2_O_2_ and ethanol response cannot be explained by lack of power, as the patterns remained robust to downsampling (Fig. S2). Overall, the number of Hog1-dependent genes for the YPS606 H_2_O_2_ response exceeded the number of Hog1-dependent genes observed for S288C NaCl response, indicating a broad non-osmotic function of Hog1 that is not observed in the laboratory strain. These results cast some doubt on Hog1’s supposed osmospecificity, which we wished to explore further.

### Hog1 displays stress-specific regulatory plasticity in wild yeast strains

To better understand how Hog1 affects gene expression under different stress conditions, we mapped stress specificity of Hog1 genotype’s effect across each strain and stress condition using ternary plots (Fig. 3, Fig. S4). We initially hypothesized that Hog1 was partially activated in wild strains during H_2_O_2_ and ethanol stress, which would lead to Hog1-dependent regulation for only the most sensitive subset of NaCl-responsive targets. However, this was not the case, as our ternary plots revealed a substantial fraction of genes that were affected in the *hog1Δ* mutant for only one stress condition, especially for YPS606 (Fig. 3A). For example, the Hog1-dependency of *AQY2* (encoding an aquaporin involved in water transport) was osmospecific–a pattern consistent with its role in osmoadpation–while the Hog1-dependency of *PTP3* (encoding a Hog1 phosphatase [83]) showed H_2_O_2_ specificity, suggesting a possible distinct feedback regulatory role during oxidative stress. There were also genes that responded to lack of *HOG1* with similar expression-change magnitudes in two stresses (most notably NaCl and H_2_O_2_), as well as a small number of genes affected by lack of *HOG1* for all stresses. For example, the known Hog1 target *GPD1* (encoding glycerol-3-phosphate dehydrogenase, a key gene necessary for fitness during osmotic stress [17]), showed similar levels of Hog1-dependency for NaCl, H_2_O_2_, and ethanol stress (Fig. 3A).

**Figure 3.**
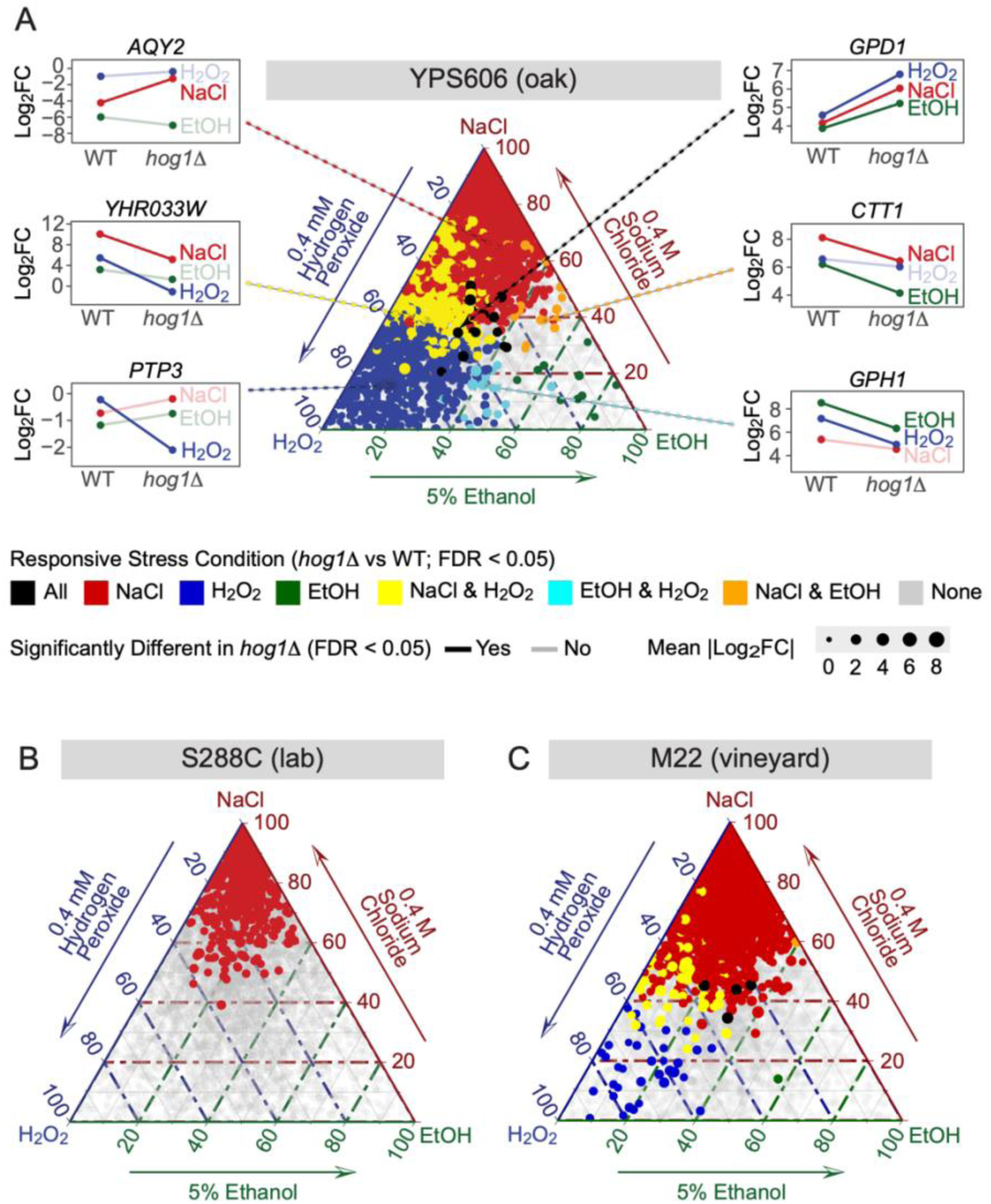
HOG1-dependent gene regulation shows stress-specific patterns in wild strains, but osmospecificity in S288C. Ternary plots show the relative magnitude of Hog1-dependent gene expression changes across the three stress conditions for YPS606 (A), S288C (B), and M22 (C). Each point represents a gene, positioned based on the proportion of its total Hog1-dependent expression change (sum of absolute log_2_ fold changes) attributable to each stress. Genes clustering near the corners exhibit stress-specific Hog1 regulation, while genes in the center display similar Hog1 dependence across all three stresses. Colors indicate which stresses the gene was significantly differentially expressed in for the *hog1Δ* vs wild-type comparison (fold change >1.5, FDR < 0.01), and point size reflects expression level (TMM-normalized logCPM values).

In contrast to YPS606, which displayed a complicated pattern of shared and stress-specific sets of Hog1-regulated genes, the S288C Hog1 regulon was highly NaCl specific (Fig. 3B). This pattern indicates that not only are genes identified as differentially expressed in S288C only identified in salt stress (consistent with Fig. 2), but also that expression of these salt-responsive Hog1 targets changes minimally in the absence of *HOG1* for the other tested stresses. M22 displayed an intermediate pattern of distribution, with fewer genes clustering along the H_2_O_2_-NaCl axes, but with greater dispersion than the S288C strain (Fig. 3C), indicating that the non-osmospecific pattern of Hog1-dependency in YPS606 is not confined to one strain background. Altogether, this analysis suggests surprising plasticity in HOG signaling in wild yeast strains responding to diverse stressors.

### Hog1 displays divergent regulation of translation and metabolism across strains and stresses

The diverse patterns of Hog1-dependency across strains and stresses suggested that different classes of genes may be Hog1-regulated under different conditions. Thus, to identify processes regulated by Hog1, we performed gene-set enrichment analysis (GO and KEGG) for each strain and stress response. Functional enrichments in NaCl stresses largely overlapped across all strains (Fig. 4), with a small number of strain-specific processes (e.g., transposition is specifically repressed by Hog1 in S288C, while genes involved in sporulation and sexual reproduction were specifically repressed by Hog1 in YPS606 and M22, respectively).

**Figure 4.**
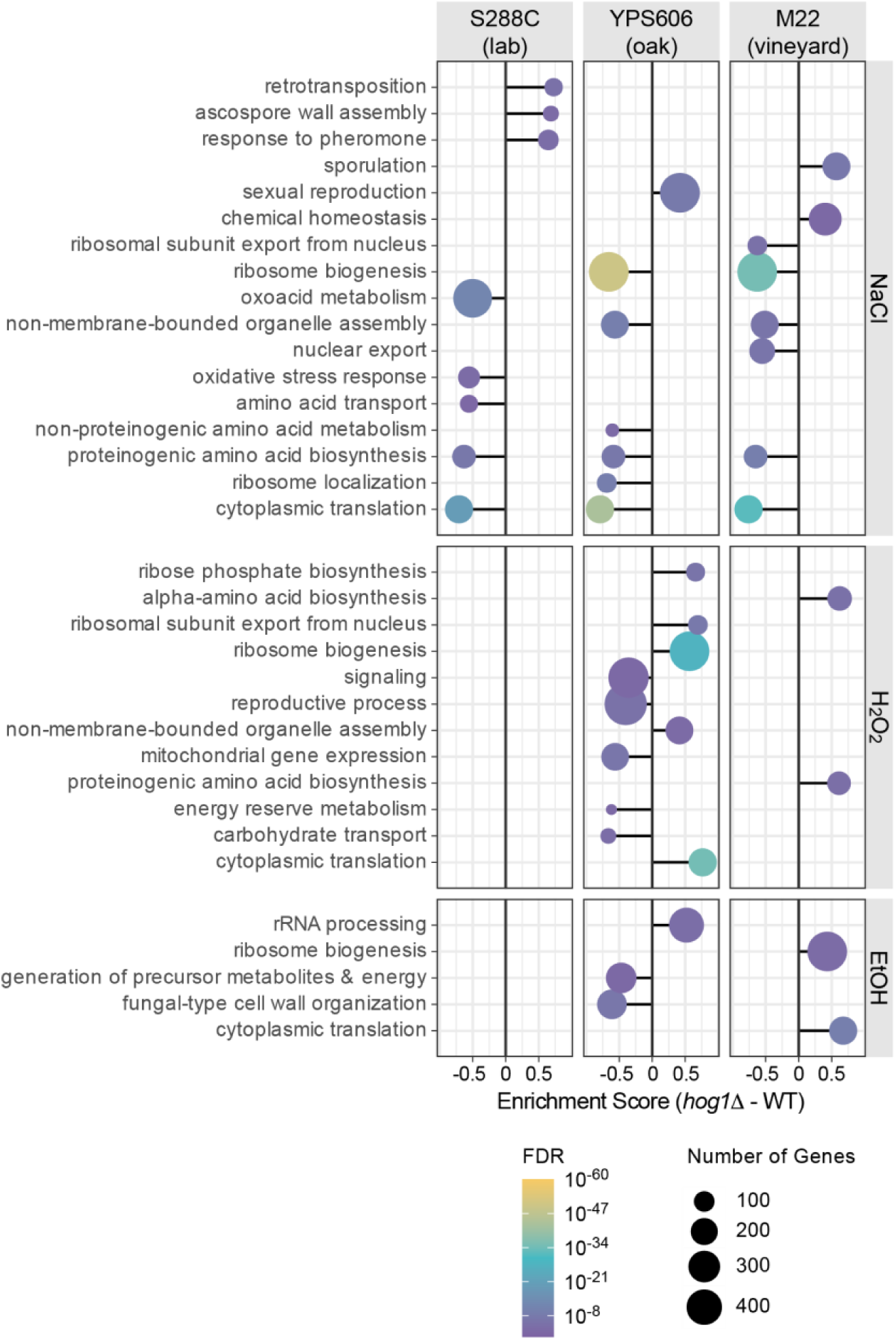
Hog1 regulates different cellular processes in wild strains responding to non-osmotic stresses. Gene set enrichment analysis (GSEA) of GO and KEGG functional categories is depicted for Hog1-dependent gene expression changes across all strain-stress combinations. Dot sizes represent the number of genes in each enriched gene set, and colors indicate significance level, according to the key. Negative enrichment scores indicate gene sets enriched among genes with lower expression in the *hog1Δ* mutant compared to the wild-type strain during stress, while positive enrichment scores indicate gene sets enriched among genes with higher expression in *hog1Δ* mutant compared to wild-type strain. No significant rank-based enrichments were identified for S288C in non-osmotic conditions. Significant terms shown for FDR < 0.01.

Consistent with differential expression analysis, we identified no significant functional enrichments for S288C (wild-type vs *hog1*Δ mutant) during H_2_O_2_ or ethanol stress. In contrast, we identified significant functional enrichments for both wild strains in both non-osmotic stress conditions. As expected from differential expression analysis, YPS606 displayed more significantly enriched terms than M22 for H_2_O_2_ and ethanol stress (Fig. 4, Table S6). Notably, the wild strains had distinct functional classes of Hog1-regulated genes for non-osmotic stresses. For example, the M22 *hog1Δ* mutant displayed higher expression of genes involved in amino acid biosynthesis during the H_2_O_2_ response, while the YPS606 *hog1Δ* mutant had lower expression of mitochondrial genes and those involved in carbohydrate transport and energy reserve metabolism. Overall, the coherent functional enrichments provide further support for a broader role for Hog1in these wild strains compared to the S288C lab strain.

One process that showed particularly striking patterns of clear, stress-specific Hog1 dependence was ribosomal protein (RP) expression. During NaCl stress, *hog1Δ* mutants in all strain backgrounds showed enhanced repression of RP transcripts (Fig. 5; Fig. S5). However, during non-osmotic stresses, this pattern was inverted in the wild strains–YPS606 and M22 *hog1Δ* mutants failed to fully repress RP transcripts during H_2_O_2_ and ethanol stress, respectively (Fig. 5). The opposing regulatory phenotypes of YPS606 and M22 *hog1Δ* mutants–with enhanced repression during osmotic stress, but defective repression during one non-osmotic stress–suggest that Hog1 differentially modulates ribosome biogenesis in a strain and stress specific manner.

**Figure 5.**
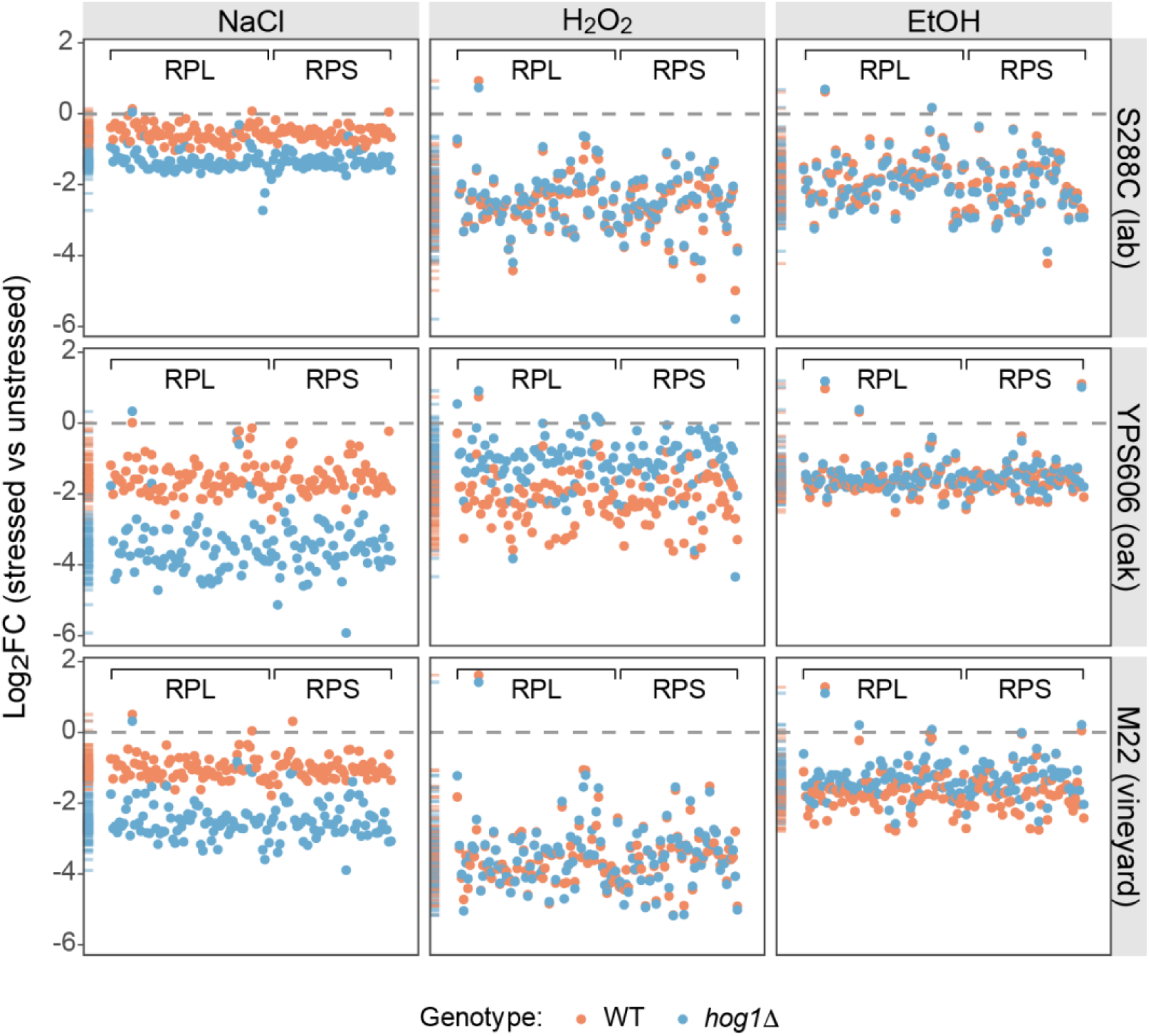
HOG1 exhibits opposing effects on ribosomal protein gene regulation between osmotic and non-osmotic stresses in wild yeast strains. Each point represents log_2_ fold change in expression for a single ribosomal protein gene (RPL/RPS) during the depicted stresses. Orange and blue depict the wild-type and *hog1*Δ stress responses, respectively. Marginal density plots on the left of each panel show the distribution of expression changes.

### Gene expression patterns in *hog1Δ* mutants correlate with acquired H_2_O_2_ resistance phenotypes

Our transcriptional profiling of *hog1Δ* mutants across strains and stresses provided an opportunity to potentially identify which Hog1-regulated genes are important for acquired stress resistance. To directly connect gene expression changes to acquired H_2_O_2_ survival, we used the machine learning approach of partial least squares regression to identify gene expression patterns that correlate with H_2_O_2_ survival (MIC*; Table S7). As expected, one of the model’s top hits (VIF>1) for genes positively correlated with H_2_O_2_ survival was *CTT1* (encoding catalase T, a key H_2_O_2_ scavenger), which we and others have shown is important for both NaCl and ethanol-induced H_2_O_2_ resistance [56,57,84] with a variance inflation factor (VIF) of 3.9. Targeted deletion of top non-*CTT1* candidates did not identify any other genes with large effects on acquired H_2_O_2_ resistance (Fig. S6).

However, *CTT1* expression could not explain all phenotypic differences observed (Fig. 6). For example, the YPS606 *hog1Δ* mutant failed to acquire H_2_O_2_ resistance following H_2_O_2_ pretreatment, yet *CTT1* expression was unaffected. Instead, the lack of acquisition better correlated with defective RP transcript repression in the YPS606 *hog1Δ* mutant during H_2_O_2_ stress (Fig. 5). Similarly, the M22 *hog1Δ* mutant displayed defective ethanol-induced acquired H_2_O_2_ resistance without a large decrease in *CTT1* expression, but again with defective RP transcript repression (Fig. 5).

**Figure 6.**
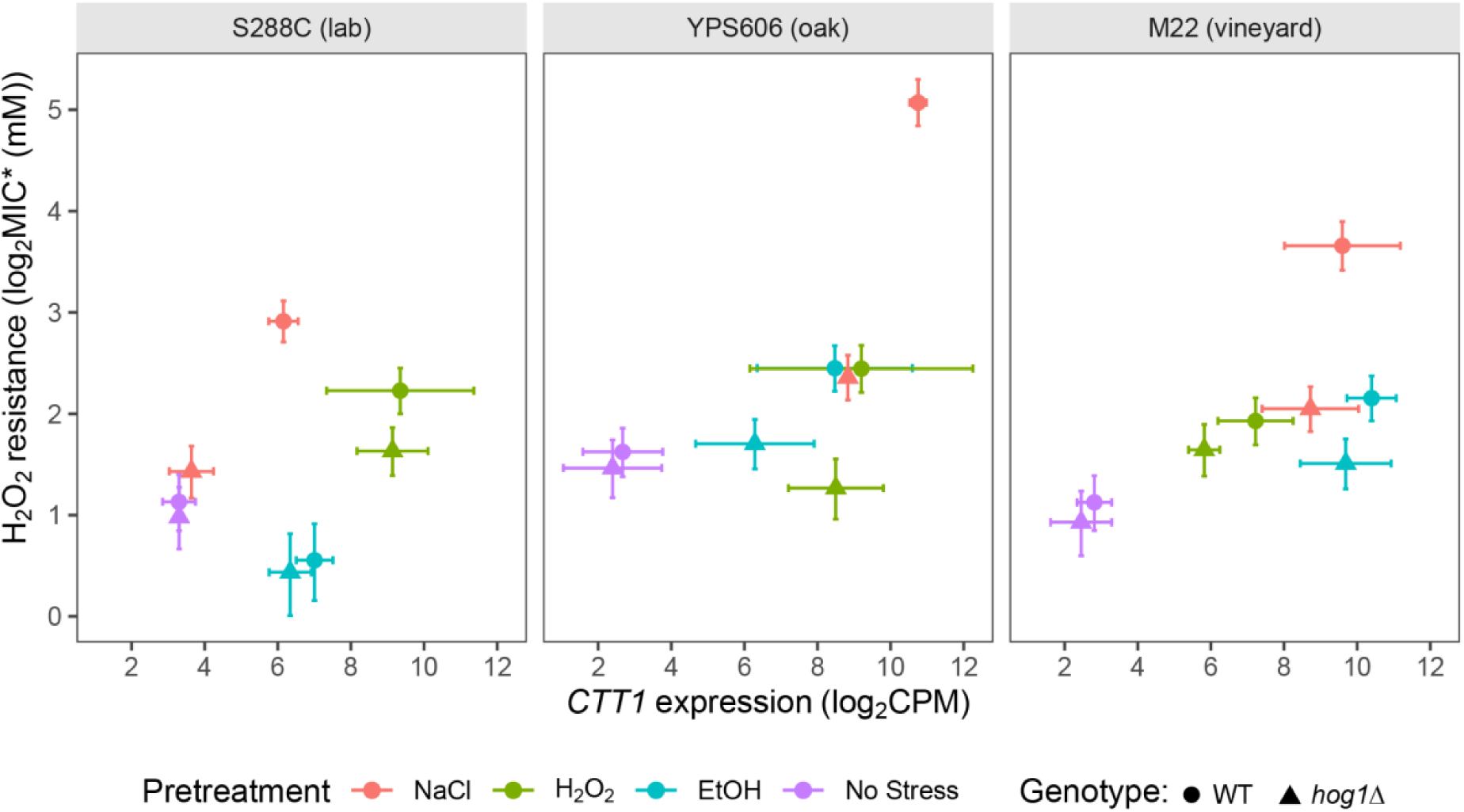
Variation in *CTT1* induction alone is insufficient to explain HOG1-dependent acquired H₂O₂ resistance. Each panel depicts *CTT1* expression levels (TMM normalized log_2_CPM values) vs acquired H₂O₂ resistance (MIC*) for wild-type (circles) and *hog1Δ* (triangles), with colors denoting stress pretreatment. Error bars denote estimated 95% confidence intervals.

There is accumulating evidence that RP repression during stress is important for redirecting translational capacity towards induced mRNAs [85,86], suggesting that defective RP repression in wild strain *hog1Δ* mutants may limit the translation of *CTT1* transcript. Notably, defective RP repression in wild strain *hog1Δ* mutants did not result in growth defects during growth on mild H_2_O_2_ or ethanol stress conditions (Fig. S7), suggesting that if defective RP repression operates at the level of acquired stress resistance rather than proliferation during stress. Altogether, these results suggest that while *CTT1* is a major effector of acquired H_2_O_2_ resistance, Hog1-dependent regulation of translational programs is likely an important strain-specific component of acquired stress resistance.

### Transcription factor target enrichments reveal context-dependent Hog1 regulation

The diversity and often altered directionality of Hog1-regulated genes and processes during non-osmotic stresses in the wild strains suggested that Hog1 may regulate different transcription factors in a strain and stress-specific manner. To identify potential transcription factors, we performed transcription factor target enrichment using the TFLink database [75] (Table S8).

As expected, the canonical osmotic stress Hog1 targets Sko1 and Hot1 showed significant target enrichment under NaCl stress in all three strains. Beyond these canonical targets, a large number of additional transcription factors were enriched during NaCl stress (91 in S288C, 101 in M22, and 114 in YPS606), likely reflecting substantial indirect effects of Hog1 on the transcriptional network. In contrast, S288C showed minimal transcription factor target enrichment for non-osmotic stresses, with only Msn2 and Rlm1 being significantly enriched during H_2_O_2_stress. YPS606 displayed substantially more transcription factor target enrichments during non-osmotic stresses, with 83 and 77 significant enrichments for ethanol and H_2_O_2_ stresses, respectively. M22 showed significant target enrichments for Msn2, Msn4, and Ifh1 during ethanol stress, and Msn4 and Met28 for H_2_O_2_stress. These patterns mirror the levels of differential expression results, with wild strains displaying network complexity during non-osmotic stresses, and S288C showing minimal Hog1-dependent regulation.

At least one of the general stress response transcription factor paralogs Msn2/4 was identified as significantly enriched in all strain-stress combinations where any transcription factor enrichment was detected. However, the directionality of the enrichment was opposite between osmotic and non-osmotic stress conditions for all strains. This manifested as higher expression for Msn2/4 targets in the *hog1*Δ mutants for osmotic stress and lower expression of a partially overlapping set of targets in *hog1*Δ mutants for non-osmotic stresses (Fig. S8). In wild-type cells, many Msn2/4 targets were not induced during osmotic stress, but exhibited strong induction in the *hog1*Δ mutant, indicating novel induction by Msn2/4 in the absence of Hog1.

We were particularly interested in identifying possible transcription factors responsible for the divergent patterns of RP expression observed in wild strain *hog1*Δ mutants across stresses. Previous research has suggested a connection between Dot6/Tod6 regulators of translational activity [87], but these TFs were not identified as enriched in this analysis, suggesting alternate mechanisms for stress-specific activity. One candidate that emerged as enriched among Hog1-regulated genes was Ifh1. Ifh1 is a known activator of RP gene expression, and its enrichment parallels the pattern of divergent RP gene expression observed across wild strains and stresses. This raises the possibility that Hog1 modulates RP transcription either directly or indirectly via Ifh1.

We previously tested several transcription factors identified in the enrichment analysis above for causal roles in acquired H_2_O_2_ resistance in the YPS606 background [58]. For NaCl-induced cross protection, none of Msn2/4, Sko1, or Hot1 were necessary, and *CTT1* induction was observed in the deletion mutants, consistent with a dominant role for *CTT1* in acquired H_2_O_2_ resistance. In contrast, Msn2/4 were necessary for ethanol-induced acquired H_2_O_2_ resistance, and *CTT1* transcript levels were dramatically reduced in the *msn2/4ΔΔ* strain during ethanol stress. Based on the expression levels of *CTT1* in *hog1Δ* mutants in wild strains under non-osmotic stresses (Fig. 6), it seems likely that at least for non-osmotic stresses, Hog1-dependency for cross protection is not due to effects on Msn2/4, Sko1, or Hot1 targets.

### Hog1 likely regulates non-osmotic stress responses non-canonically in wild strains

The dramatic differences in Hog1-dependent gene expression changes in wild strains responding to non-osmotic stresses could be the result of differences in Hog1activation itself. Canonically, during osmotic stress Hog1 is phosphorylated by Pbs2 via activation of two main branches (Sln1 and Sho1). Hog1∼P can phosphorylate cytoplasmic targets, but it largely affects gene expression by translocating to the nucleus and phosphorylating stress-responsive transcription factors.

We thus first examined via western blot whether Hog1 was phosphorylated in each strain background during different stress exposures. In response to NaCl exposure, Hog1 showed robust phosphorylation (Fig. 7A) within 5 minutes of stress treatment for each strain, as has been previously reported [27,40]. However, even in the wild strains, no phosphorylation was observed for H_2_O_2_ and ethanol stresses at 5 minutes of stress exposure. Likewise, Hog1-GFP accumulated in the nucleus within 5 minutes of NaCl exposure for all strains (Fig. 8), but no nuclear localization was detected for the non-osmotic stresses. To test whether Hog1 activation dynamics differed in response to H_2_O_2_ and ethanol in the wild strains, we further examined Hog1 phosphorylation and nuclear localization at later timepoints. We did observe delayed but strong phosphorylation of Hog1 at 30 minutes post-H_2_O_2_ exposure in YPS606 but not S288C (Fig 7B). However, we still did not observe any Hog1∼P in response to ethanol stress at 30 minutes of exposure in either YPS606 or S288C, though we hypothesize that low levels of Hog1 phosphorylation do occur during ethanol stress in the wild strains, but were below our detection limit. However, despite observing delayed Hog1∼P during H_2_O_2_ stress, we still did not detect Hog1-GFP nuclear localization above unstressed background between 5 and 60 minutes of H_2_O_2_ stress in YPS606 (Fig. S9 and Supplementary Materials). Notably, Hog1 effects on transcription in the wild strains were observed at 30 minutes post-stress exposure for H_2_O_2_ and ethanol stress, and the regulons were largely distinct. Those observations, combined with the delayed phosphorylation of Hog1 during H_2_O_2_ stress, and the lack of detectable nuclear localization for both H_2_O_2_ and ethanol stresses, suggest that Hog1 is regulating transcription during non-osmotic stresses in wild strains non-canonically, likely through cytoplasmic targets (see Discussion).

**Figure 7.**
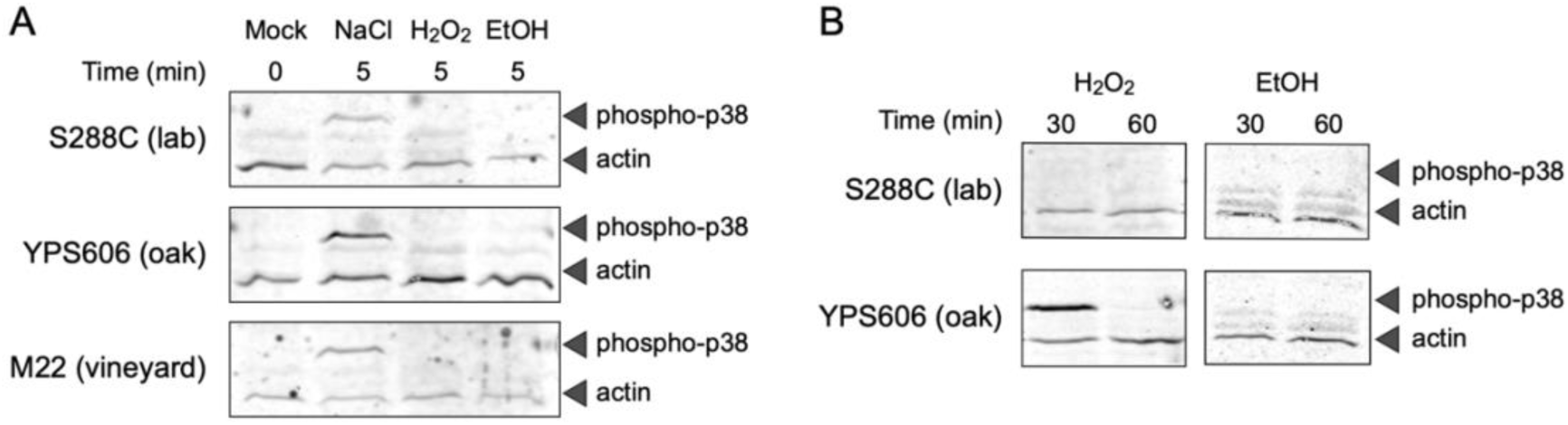
Hog1 phosphorylation is rapid during NaCl stress in all strains but is delayed during H₂O₂ stress in YPS606. Western blot analysis was performed using a cross-species Hog1∼P antibody (anti-phospho-p38 MAPK) and anti-actin as a loading control. Hog1 phosphorylation was measured at 5-min post stress for all strains (A), and at 30- or 60-min post-EtOH and H₂O₂ for S288C and YPS606 (B). Representative western blots are depicted, and all replicates can be found in the Supplementary Materials.

**Figure 8.**
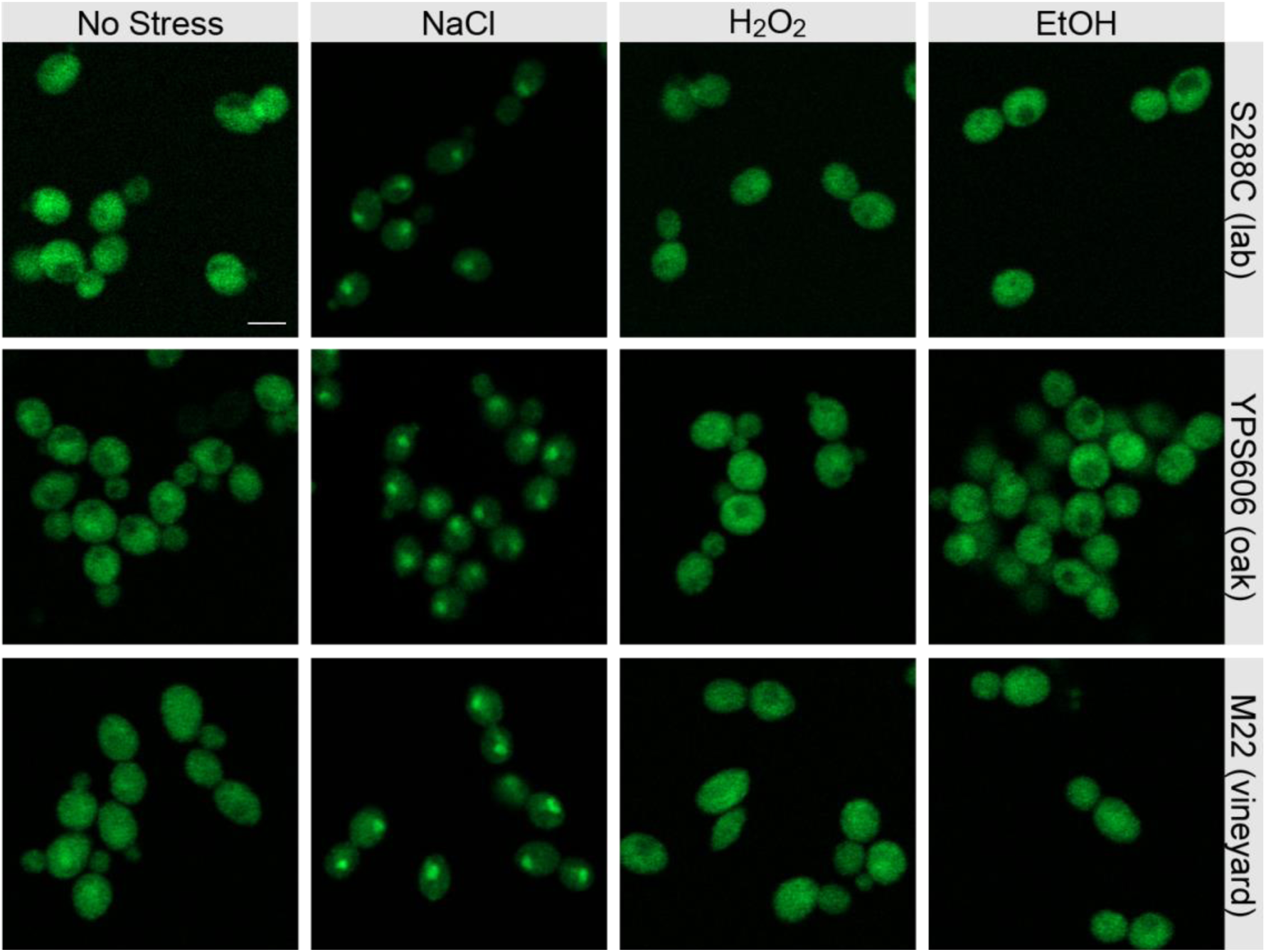
Across all strains, Hog1-GFP rapidly translocates to the nucleus during NaCl stress but remains cytoplasmic during H₂O₂ and ethanol stress. Each panel depicts live-cell imaging of Hog1-GFP following 5 min of stress exposure. Scale bar: 10 microns.

## Discussion

This study suggests that the prevailing view of Hog1 as largely osmo-specialist in *S. cerevisiae* is incomplete and has been shaped by disproportionate studies performed in laboratory strains that are genetic and phenotypic outliers. By analyzing stress defense phenotypes dependent on stress-activated gene expression changes and transcriptomic analyses across diverse strain backgrounds, we demonstrate that Hog1 plays a significant role in non-osmotic stress responses in wild yeast strains.

This conclusion is supported by three main findings. First, deletion of *HOG1* reduced acquired H_2_O_2_ resistance following at least one non-osmotic pretreatment in all wild strains tested. Second, *hog1Δ* mutants in wild strains displayed dozens to hundreds of differentially expressed genes during non-osmotic stress, while no expression changes were observed in S288C under the same conditions. Third, activation of Hog1 via phosphorylation in wild strains during non-osmotic stress exhibits distinct kinetics compared to the canonical osmotic response, and no nuclear Hog1 was observed, suggesting a unique cytoplasmic role. Together, these findings indicate strain-dependent plasticity in HOG signaling, driven by non-canonical HOG activation during non-osmotic stresses rather than attenuated canonical signaling.

### Why are *hog1Δ* mutants in wild yeast strains generally unable to fully acquire futher H_2_O_2_ resistance when pretreated with osmotic and non-osmotic stresses?

We previously showed that *CTT1* encoding catalase plays a large role in acquired H_2_O_2_ resistance [57,58]. However, *CTT1* mRNA levels in *hog1Δ* mutants were often unaffected by the deletion, particularly during non-osmotic stresses, suggesting that effects on *CTT1* transcriptional induction alone cannot explain the observed phenotypes. We examined deletions of other genes whose expression in *hog1Δ* mutants most strongly correlated with acquired H_2_O_2_ resistance, and all were still able to acquire high levels of further H_2_O_2_ resistance (Fig. S6). This leads us to propose an alternative explanation, where the predominant link between Hog1-dependent gene expression and survival appears to be regulation of translational machinery. Under NaCl stress, exaggerated repression of ribosomal protein (RP) genes in *hog1Δ* mutants is consistent with Hog1-mediated maintenance of translation, possibly through phosphorylation of Rck2 or modulation of Dot6/Tod6 activity [16,45,87,88]. In contrast, under H_2_O_2_ stress in the YPS606 oak strain, failure of *hog1Δ* mutants to fully repress RP expression coincided with loss of acquired H_2_O_2_ resistance. This bidirectionality–enhanced repression during osmotic stress, but defective repression during oxidative stress–suggests that Hog1 is not merely activating osmotic programming in response to non-osmotic stresses. Rather, Hog1 appears to actively regulate translational capacity in a stress-specific manner in the wild strains.

The enrichment of targets for the RP regulator Ifh1 and general stress response factors Msn2/4 in the Hog1-dependent gene sets of wild strains further supports a model where Hog1 acts as a coordinating regulatory hub for balancing translation of stress-induced transcripts, a role that may be largely lost in S288C. Two mechanisms potentially explain why *hog1Δ* mutants exhibit defective acquired H_2_O_2_ resistance: (1) failure to repress RP/ribosome biogenesis (RiBi) gene expression reduces the translation of stress-induced transcripts [16], and (2) inappropriate post-transcriptional regulation of H_2_O_2_ defenses, perhaps involving stress granule sequestration of stress-induced transcripts like *CTT1*, thus reducing their translation [39,89,90].

The mechanism underlying divergent Hog1 function appears to lie in upstream signaling dynamics and subcellular localization. Our data are consistent with a model in which non-osmotic stresses such as H_2_O_2_ activate Hog1 through distinct mechanisms, such as phosphatase inhibition or stress granule scaffolding as observed by Lee and colleagues [39,89,90], rather than robust kinase cascade activation and nuclear localization as seen in osmostress. The delayed yet high-amplitude phosphorylation signal observed in YPS606 without detectable nuclear accumulation points toward a primarily cytoplasmic role for Hog1 in this context. This could involve direct regulation of translational machinery components via intermediate regulators like the MAPK-activated kinase Rck2 [45,88], or modulation of channel proteins like Fps1, or via to this point undescribed targets

While we cannot rule out transient nuclear pulses below our detection limit, we found no evidence for Hog1 nuclear localization during non-osmotic stresses in the wild strains despite an extensive time course. Moreover, the stress-specific signatures of the regulons, with largely distinct genes being Hog1-dependent during non-osmotic vs osmotic stresses, argue against a model of proportionally weaker activation of canonical HOG signaling during non-osmotic stress in the wild strains. The regulation of highly sensitive Hog1 targets like *STL1* (Fig. 9) during non-osmotic stress could occur through indirect mechanisms or may require only minimal transient nuclear localization below our detection threshold. Overall however, our data strongly suggest that the majority of Hog1-regulated gene expression during non-osmotic stress are mediated non-canonically.

**Figure 9.**
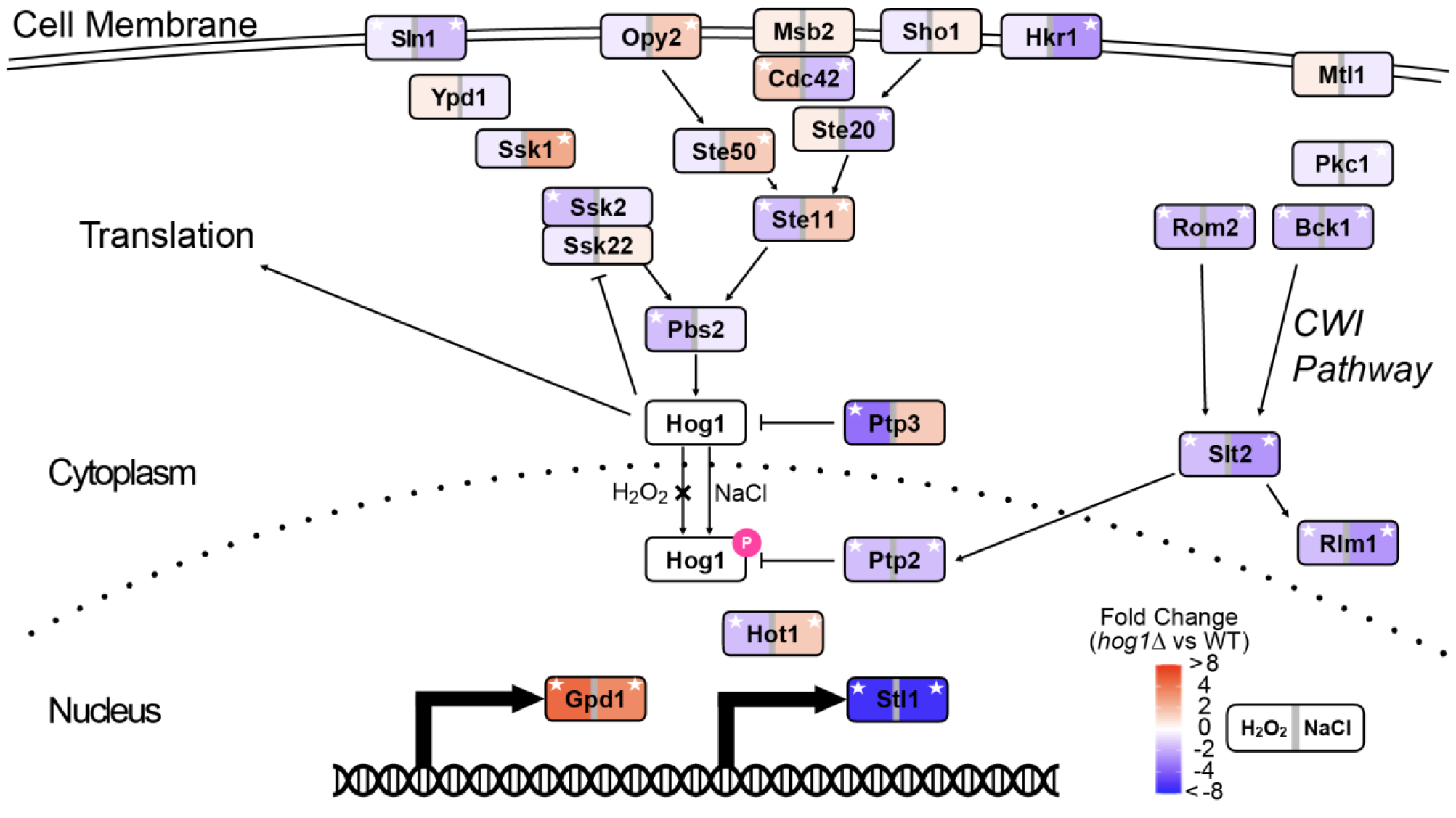
HOG pathway components show stress-specific expression patterns despite shared regulation of a subset of canonical targets. Schematic of the HOG signaling pathway with genes colored by their expression changes in the YPS606 *hog1Δ* vs wild type during H₂O₂ stress (left) or NaCl stress (right). Red denotes higher expression in the *hog1Δ* mutant vs wild type, and blue denotes lower expression, according to the key. White stars indicate significant differential expression (FDR < 0.01). Canonical HOG targets *GPD1* and *STL1* show similar Hog1-dependent regulation across both stresses, while many upstream pathway components display stress-specific patterns. CWI denotes the cell wall integrity pathway.

### MAPK regulatory plasticity and its evolutionary implications

Our results fit within a larger body of literature on HOG signaling, while clarifying why the breadth of HOG function in S288C appears narrow. The absence of ethanol-induced cross-protection against H_2_O_2_ in S288C masks any *hog1Δ* effect. Meanwhile, known genetic idiosyncrasies such as a known mutation in *HAP1* that affects catalase expression, underscore the extent to which laboratory adaptation can alter physiological responses, whether through masking of ancestral functions or selecting for altered pathway wiring through interconnected signaling pathways [91–94]. By decoupling baseline stress sensitivity from acquired stress resistance, we revealed divergence in Hog1-dependent stress responses regardless of absolute levels of resistance. This strongly argues that inference about a pathway’s functional scope within a species needs to be determined across diverse genetic backgrounds to avoid misleading, strain-specific conclusions.

The pattern we observe—osmospecificity in S288C but broader stress responsiveness in wild strains—strongly suggests that laboratory adaptation actively restricted HOG pathway function rather than multiple wild strains having evolved enhanced responsiveness. Multiple lines of evidence support this interpretation. First, wild strains from diverse ecological niches (oak soil, vineyard, distillery) all independently retain non-osmotic Hog1 functions, making convergent evolution unlikely. Second, the pathogenic yeast *C. albicans* also displays broadly stress-responsive Hog1 function [32–34], suggesting this may be ancestral in the *Saccharomyces* lineage. Third, even in S288C there is evidence that Hog1 does play a role in responding to other stresses beyond osmotic [25,28,35–43,50], though to the best of our knowledge, gene expression has not been examined until this work. Notably, we also observed fewer Hog1-regulated genes during osmotic stress in S288C compared to the wild strains, suggesting that even canonical HOG signaling is somewhat dampened in S288C at the NaCl concentrations used. S288C is notably more salt resistant than the wild strains tested, owing to increased copy numbers of the *ENA* genes encoding sodium efflux pumps [51,95], so the reduced canonical Hog1 signaling could be due to a decreased perception of the level of stress. This is unlikely the case for H_2_O_2_ and ethanol stresses, where S288C is not more resistant than the wild strains [52]. Overall, these observations indicate that broad stress responsiveness likely represents the ancestral state of HOG signaling in *S. cerevisiae*, with S288C having experienced a derived loss of function, potentially through the accumulation of multiple domestication-associated mutations that collectively dampened both canonical and non-canonical HOG signaling. We also notably observe variation in HOG signaling across the wild strains, suggesting that expansion or contraction of MAPK signaling inputs may be relatively easy to achieve evolutionarily.

### Limitations

This work has several limitations that frame future directions. Our transcriptomic analysis used single timepoints chosen to capture peak responses, which may obscure finer kinetic differences in Hog1 activation or downstream effects. Future work using ribosomal profiling and phosphoproteomics across strains and stresses will be particularly informative for dissecting post-transcriptional and post-translational components of Hog1-mediated regulation during non-osmotic stresses. Furthermore, while our expression-phenotype correlation identified *CTT1* and other known stress-responsive genes as among top predictors of cross-protection to peroxide, the distinction between correlation and causation requires systematic genetic perturbation across different backgrounds. We performed targeted deletions of high-ranking candidates beyond *CTT1*, but found no additional single genes with large effects on acquired H_2_O_2_ resistance. This suggests that the effects of Hog1 on acquired H_2_O_2_ resistance either emerge from coordinated changes across multiple genes, or via translational regulation of key stress protectants like Ctt1.

## Conclusion

In wild *S. cerevisiae* strains, Hog1 contributes to non-osmotic cross-protection and reshapes the transcriptomic responses to H_2_O_2_ and ethanol stress without detectable nuclear localization, pointing to translational control as an integral component of non-canonical HOG activity. In contrast, Hog1 was osmospecific in our commonly-used S288C lab strain, suggesting that laboratory adaptation has concealed pleiotropic roles for Hog1 that are readily apparent in wild strains. Hog1 appears to tune translational capacity in a stress and strain-specific manner [96]. Given the parallels between Hog1 and p38 kinases in higher eukaryotes [7,97], our findings suggest that such context-specific plasticity is likely an ancient, conserved property of stress-activated MAPK pathways.

## Acknowledgments

This work was supported in part by National Science Foundation grant MCB-1941824 (JAL), 5R01GM147372 (ACP), and the Arkansas Biosciences Institute (Arkansas Settlement Proceeds Act of 2000) (ACP). This research used resources available through the Arkansas High Performance Computing Center, which is funded through multiple National Science Foundation grants and the Arkansas Economic Development Commission.

## Supplemental Figures

**Figure S1.**
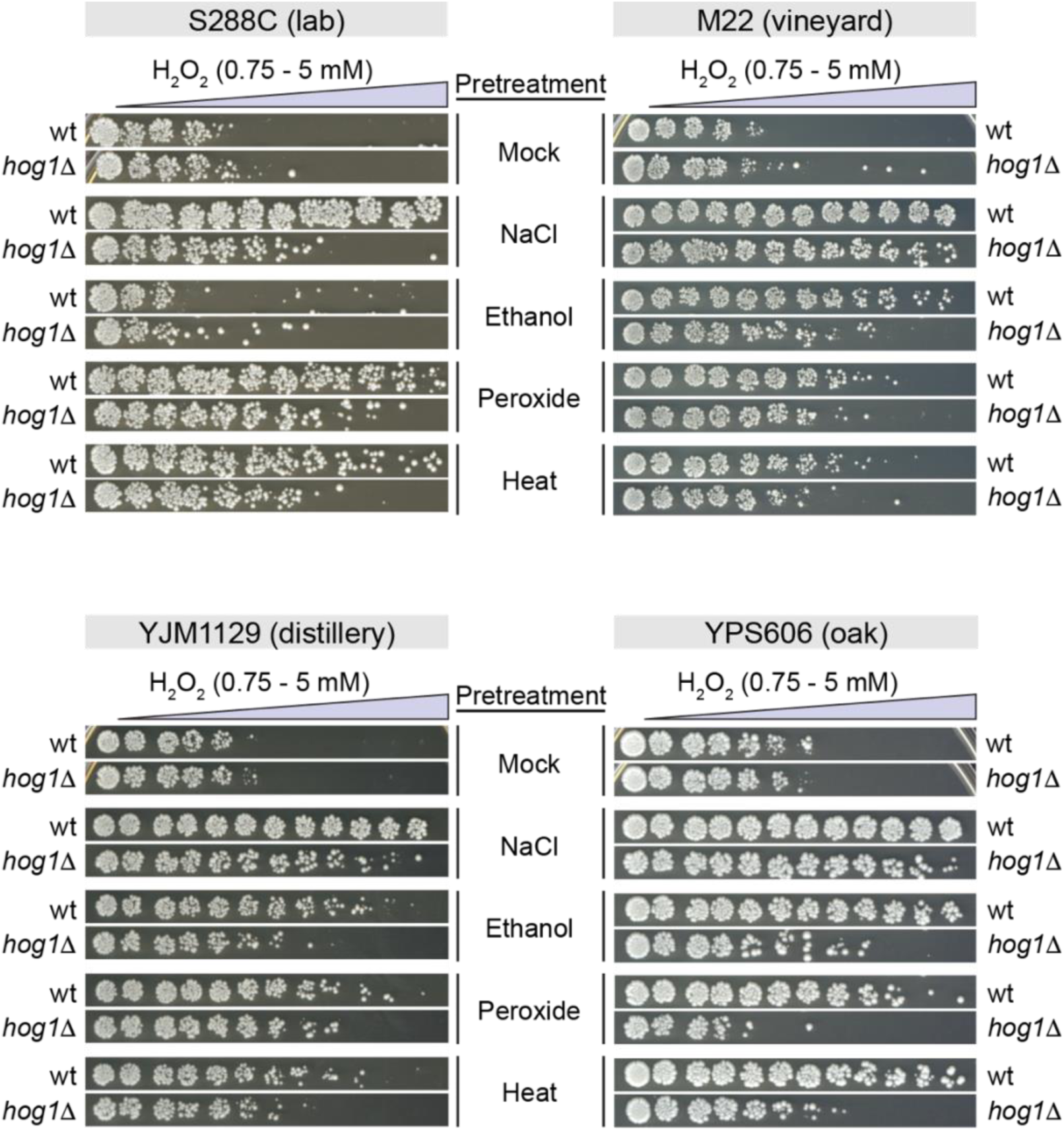
Representative H_2_O_2_ cross protection assays for all strains and conditions depicted in Figure 1.

**Figure S2.**
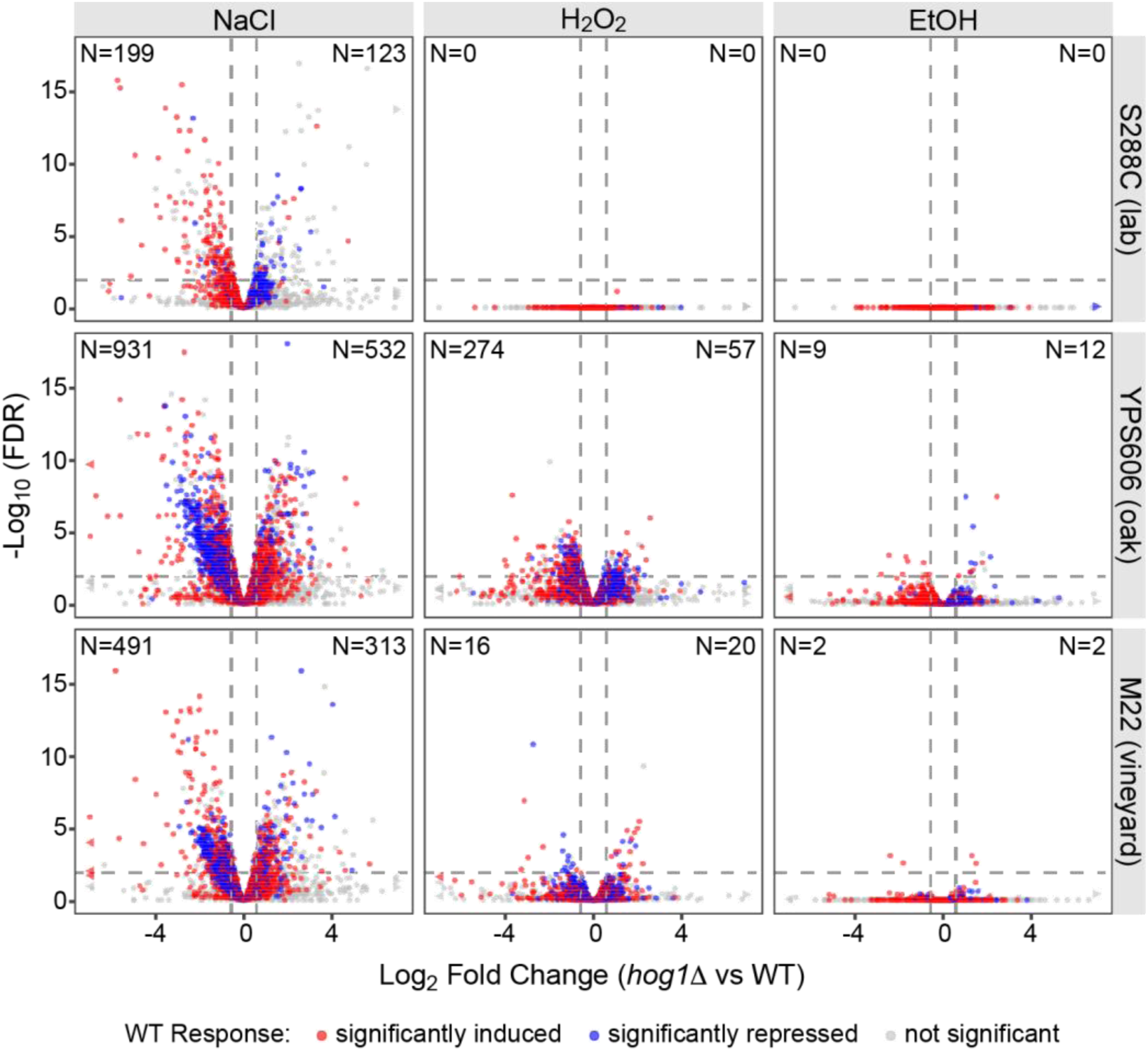
Hog1-dependent differential expression during non-osmotic stresses in wild strains is robust to subsampling and thus not due to differences in statistical power. Subsampling was performed by reducing the number of samples in wild strains to 3 (equal to S288C), and reads were subsampled down to the sample with lowest read counts. Differential expression analysis was performed in edgeR using the exact same parameters as for the full dataset. Volcano plots were performed as in Figure 2.

**Figure S3.**
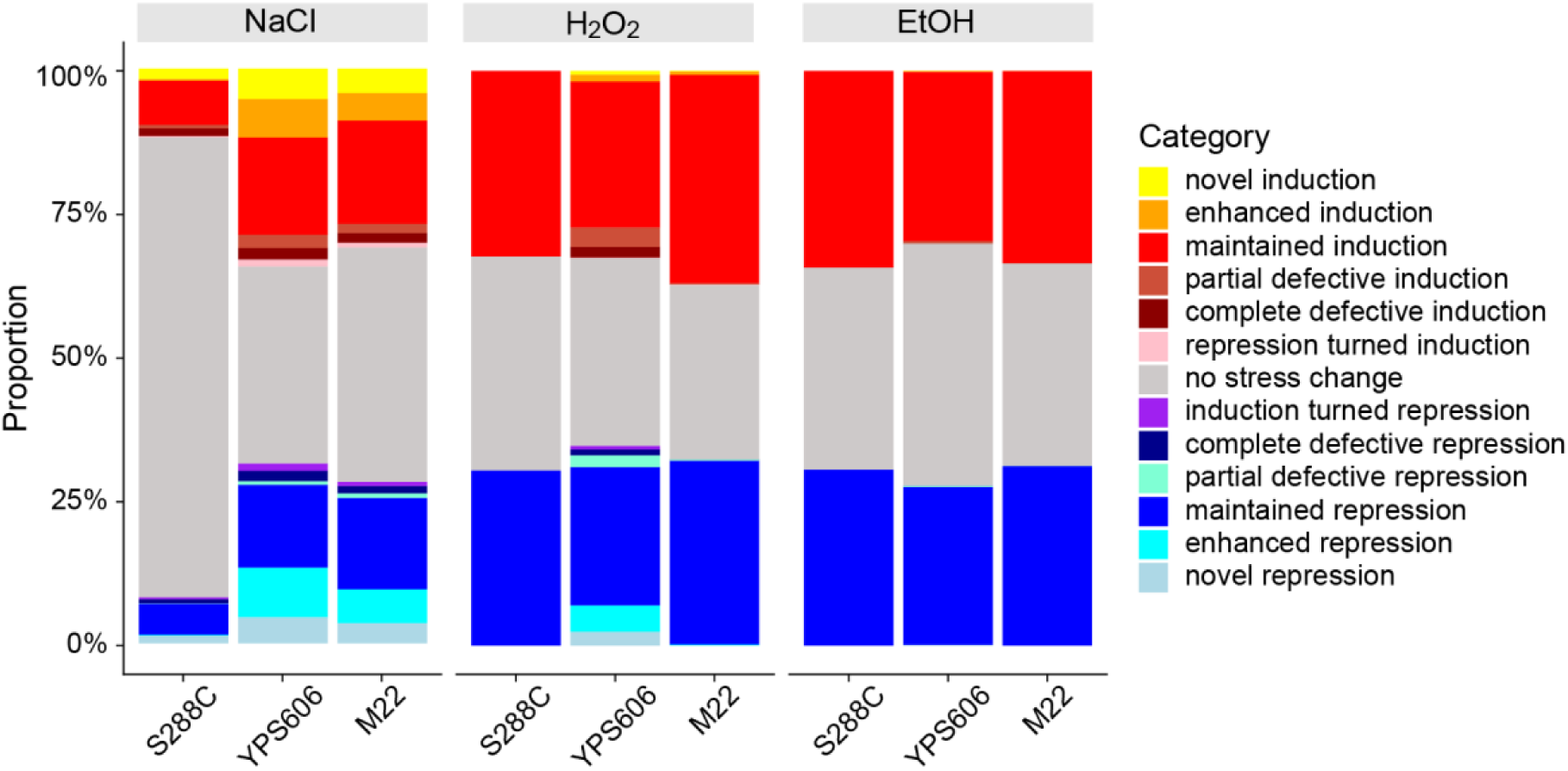
Strain-specific variation in classes of Hog1-regulated genes. The bars depict proportions of each class of genes. Genes unaffected by Hog1 fall into three major classes: no change in expression during stress (grey), induced during stress but unaffected by lack of Hog1(red), and repressed during stress but unaffected by lack of Hog1 (blue). All other classifications for Hog1-regulated genes are noted in the key.

**Figure S4.**
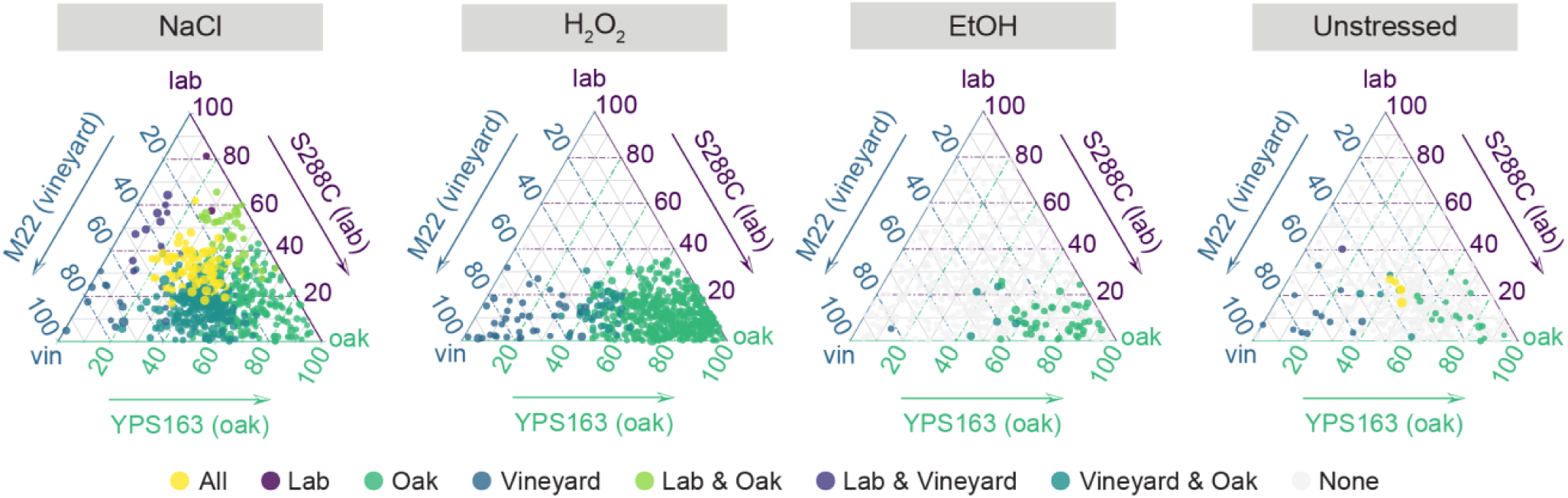
Shared and strain-specific Hog1 dependency for osmotic and non-osmotic stresses. Ternary plots show the relative magnitude of Hog1-dependent gene expression changes across all three strains as per Figure 3.

**Figure S5.**
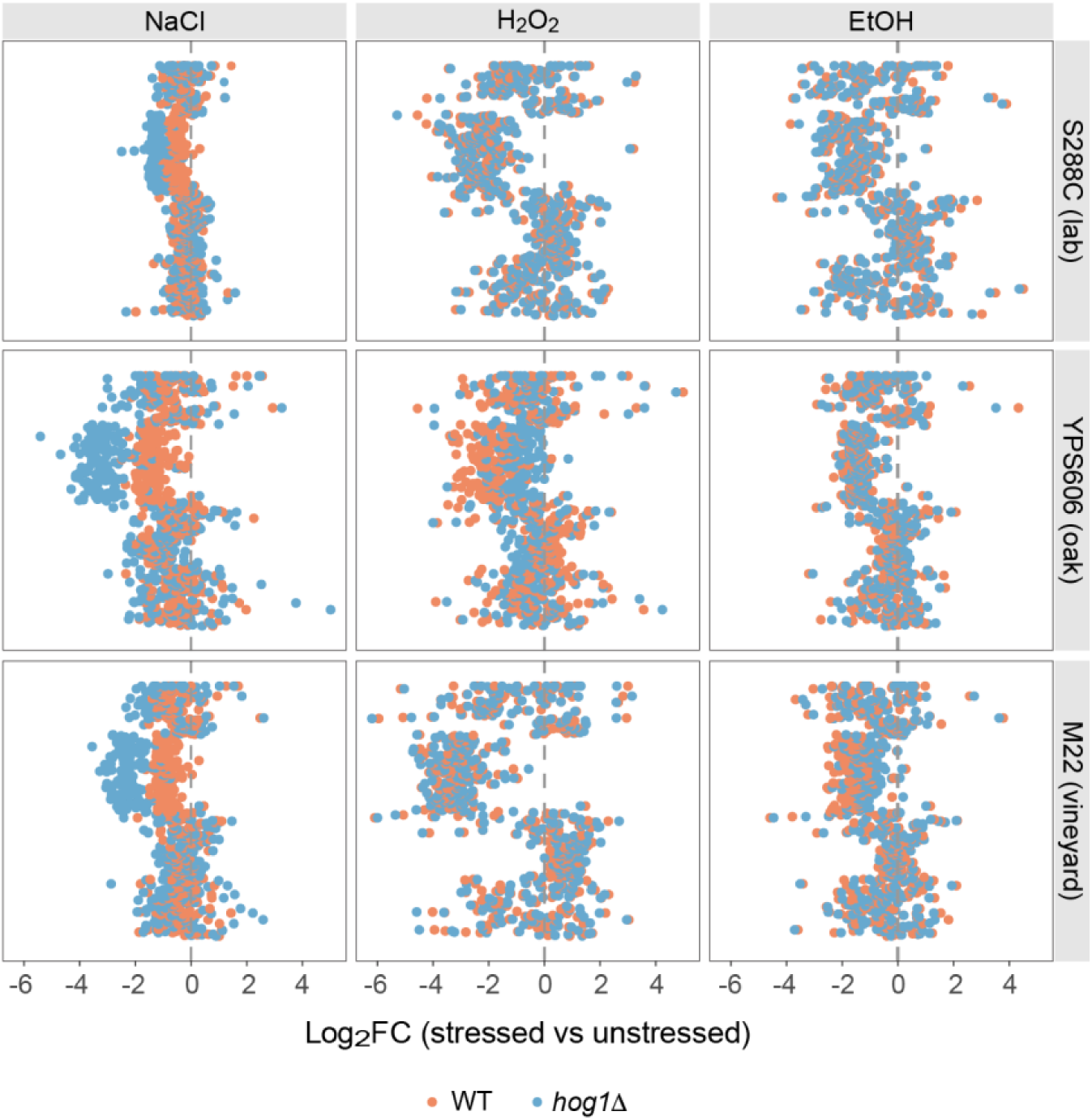
Hog1 specifically exhibits opposing effects on ribosomal protein (RP) gene regulation. All transcripts annotated as belonging to the GO term translation were sorted by gene name, which largely sorts the major classes of RP genes and ribosome biogenesis genes (Ribi).

**Figure S6.**
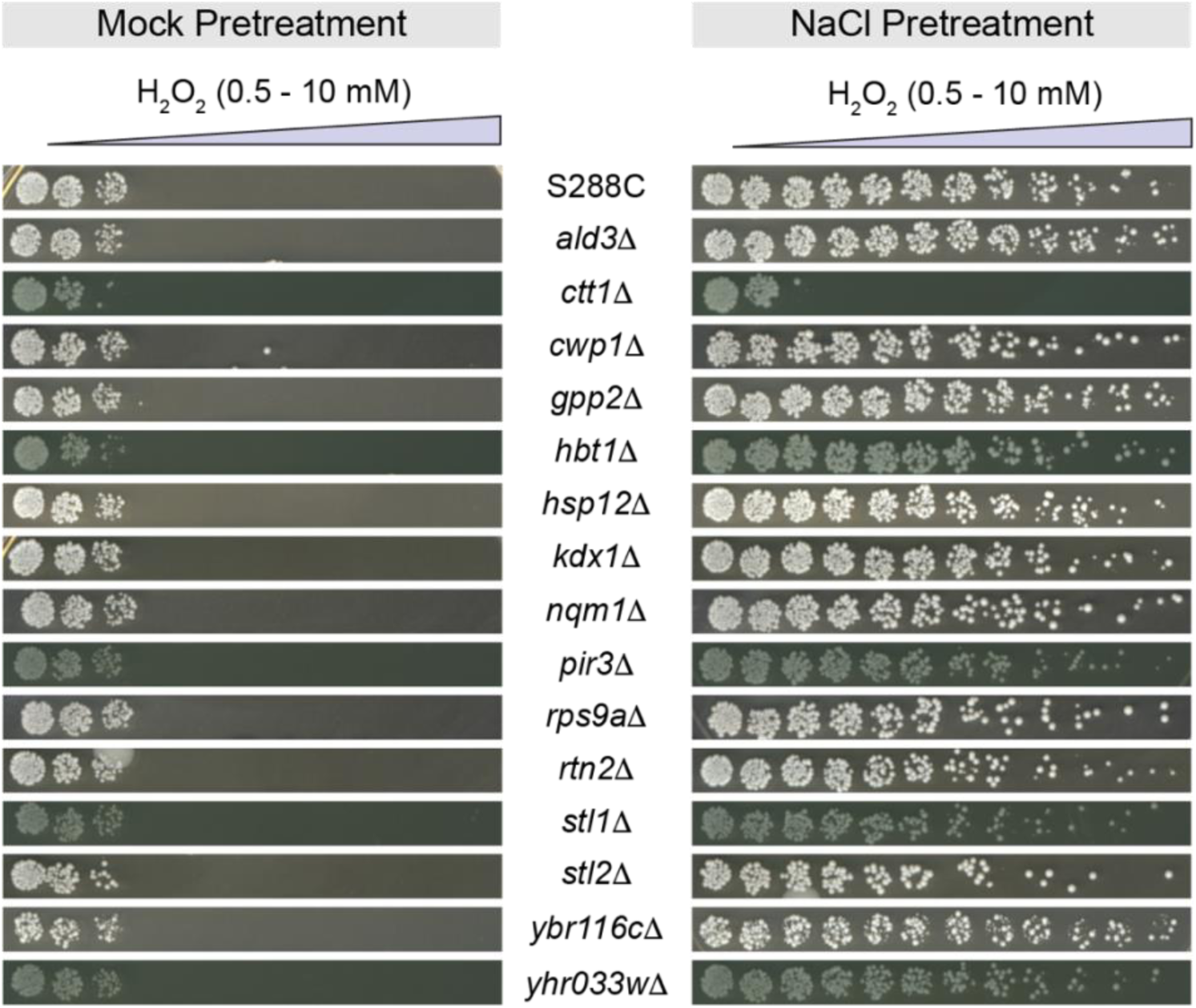
NaCl-induced cross protection phenotypes for mutants in genes implicated by correlation analysis. Strains from the YKO collection (S288C background) were chosen based on high expression-phenotype (MIC*) correlation.

**Figure S7.**
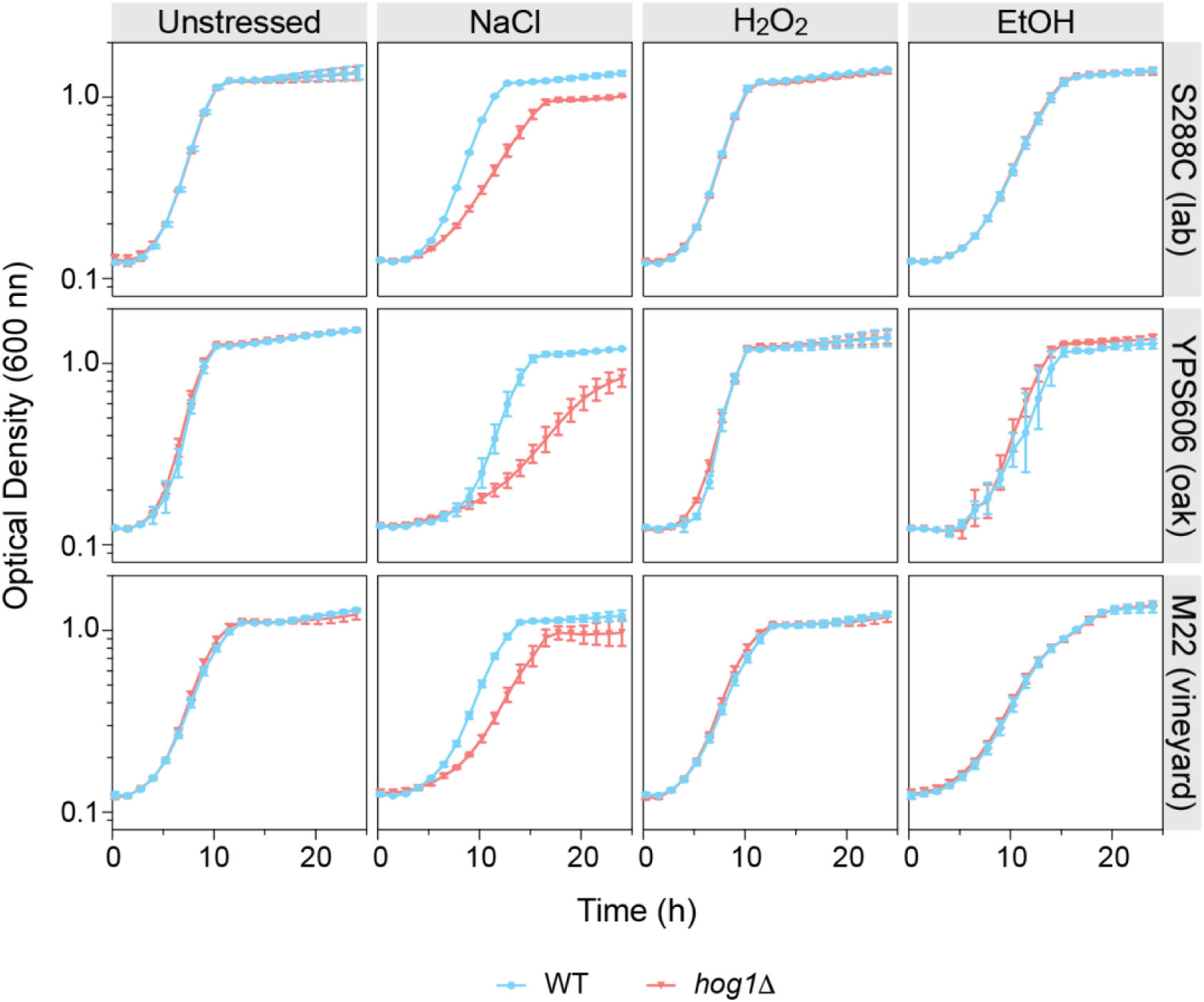
Growth analysis of wild-type and *hog1Δ* mutants under stress. Cells were grown in YPD (unstressed) or the following stress concentrations: 0.4 mM H_2_O_2_, 5% ethanol, or 0.4M NaCl. Error bars indicate the standard deviation of the mean of biological triplicates.

**Figure S8.**
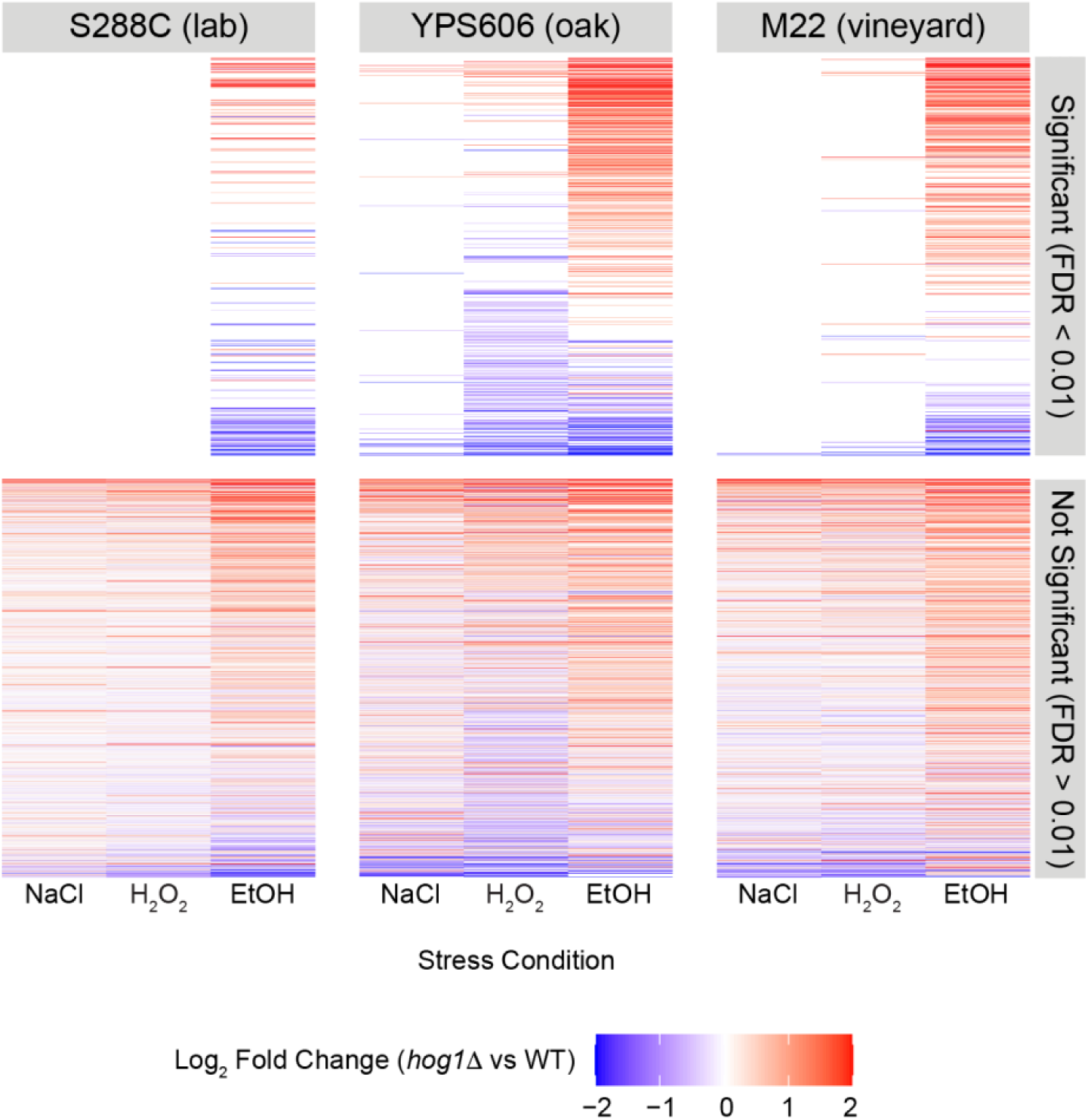
Expression patterns for Msn2/4 targets differ by stress. The heatmap depicts log_2_ fold changes in the stress response of *hog1Δ* vs WT strains in each stress condition for known Msn2/4 target genes (from the TFLink Database). The top panel depicts targets that were differentially expressed in at least one strain in the *hog1Δ* vs WT stress response comparison, while the bottom panel depicts genes that were not significantly differentially expressed.

**Figure S9.**
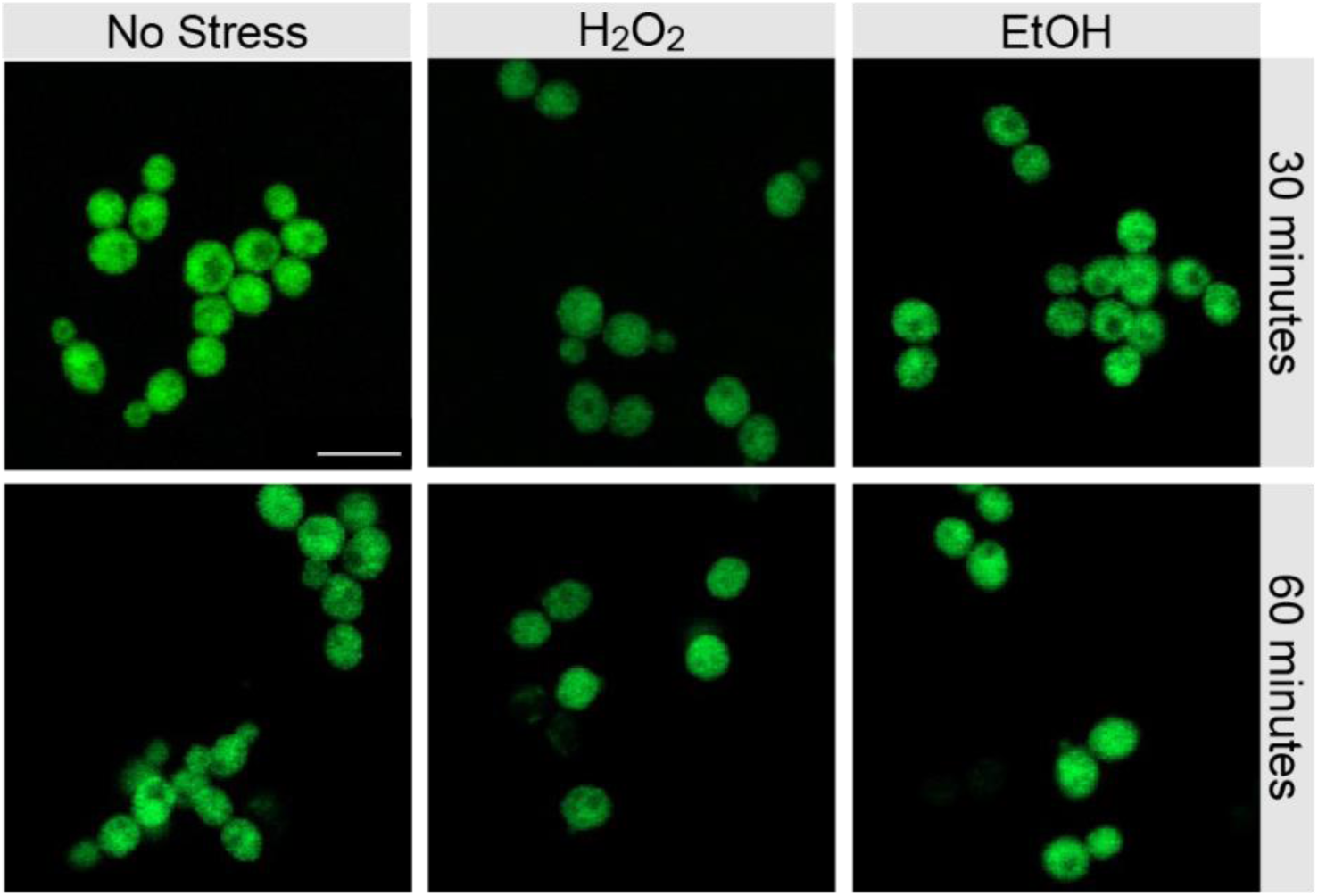
Hog1-GFP remains cytoplasmic during H₂O₂ and ethanol stress at 30 and 60 minutes in YPS606. Each panel depicts live-cell imaging of Hog1-GFP following 30 or 60 min of stress exposure. Scale bar: 10 microns.

## Supplemental Tables

**Table S1: Strains used in the study**

**Table S2: Oligonucleotides used in the study**

**Table S3: MIC* estimates for all strain-genotype combinations**

**Table S4: ΔMIC* estimates comparing wild-type strains and *hog1Δ* mutants**

**Table S5: edgeR output for differential expression analysis**

**Table S6: Functional enrichments**

**Table S7: PLS regression coefficients for predicting MIC* from gene expression**

